# Metabolic Diversity in Human Non-Small Cell Lung Cancer Cells

**DOI:** 10.1101/561688

**Authors:** Pei-Hsuan Chen, Ling Cai, Kenneth Huffman, Chendong Yang, Jiyeon Kim, Brandon Faubert, Lindsey Boroughs, Bookyung Ko, Jessica Sudderth, Elizabeth A. McMillan, Luc Girard, Michael Peyton, Misty D. Shields, David Shames, Hyun Seok Kim, Brenda Timmons, Ikuo Sekine, Rebecca Britt, Stephanie Weber, Lauren A. Byers, John V. Heymach, Michael A. White, John D. Minna, Guanghua Xiao, Ralph J. DeBerardinis

**Affiliations:** Children’s Medical Center Research Institute at UT Southwestern Medical Center, 5323 Harry Hines Blvd, Dallas, TX, 75390, USA; Department of Cancer Biology, Dana-Farber Cancer Institute, Boston, MA 02115, USA; Quantitative Biomedical Research Center, Department of Clinical Sciences at UT Southwestern Medical Center, 5323 Harry Hines Blvd, Dallas, TX, 75390, USA; Department of Cell Biology, UTSW Medical Center, Dallas, Texas 75390, USA; Laboratory of Systems Cancer Biology, The Rockefeller University, New York, NY, 10065, USA; Department of Pharmacology, University of Texas Southwestern Medical Center, 5323 Harry Hines Blvd, Dallas, TX, 75390 USA; Hamon Center for Therapeutic Oncology, University of Texas Southwestern Medical Center, 5323 Harry Hines Blvd, Dallas, TX, 75390, USA; Department of Oncology Biomarker Development, Genentech Inc, South San Francisco, California 94080, USA; Severance Biomedical Science Institute, Brain Korea 21 Plus Project for Medical Science, Yonsei University, College of Medicine, Seoul, 120-749, Republic of Korea; El Centro College, 801 Main St, Dallas, TX 75202, USA; Department of Thoracic/Head and Neck Medical Oncology, University of Texas, MD Anderson Cancer Center, Houston, Texas, 77030, USA; Pfizer Inc. New York City, New York, 10017, USA; Department of Internal Medicine, University of Texas Southwestern Medical Center, 5323 Harry Hines Blvd, Dallas, TX, 75390 USA; Howard Hughes Medical Institute, University of Texas Southwestern Medical Center, 5323 Harry Hines Blvd, Dallas, TX, 75390 USA

**Keywords:** Non-small cell lung cancer, cancer metabolism, cell lines, ^13^C stable isotope labeling, glucose, glutamine, oncogenotypes, gene expression, protein expression, therapeutic sensitivity

## Abstract

Intermediary metabolism in cancer cells is regulated by diverse cell-autonomous processes including signal transduction and gene expression patterns arising from specific oncogenotypes and cell lineages. Although it is well established that metabolic reprogramming is a hallmark of cancer, we lack a full view of the diversity of metabolic programs in cancer cells and an unbiased assessment of the associations between metabolic pathway preferences and other cell-autonomous processes. Here we quantified over 100 metabolic features, mostly from ^13^C enrichment of molecules from central carbon metabolism, in over 80 non-small cell lung cancer (NSCLC) cell lines cultured under identical conditions. Because these cell lines were extensively annotated for oncogenotype, gene expression, protein expression and therapeutic sensitivity, the resulting database enables the user to uncover new relationships between metabolism and these orthogonal processes.

## Introduction

Malignant cells reprogram their metabolism to support various steps in cancer progression. Because metabolic reprogramming is a hallmark of cancer, targeting metabolism is considered to be a potential therapeutic strategy (Hanahan and Weinberg, 2011; Tennant et al., 2010). This concept dates back to the 1920s, when Otto Warburg observed high rates of glucose uptake and lactate secretion (the Warburg effect) in tumor tissue from mice and humans (Koppenol et al., 2011). The copious lactate secretion, even in the presence of sufficient oxygen to oxidize glucose completely, was originally interpreted as evidence that suppressed mitochondrial metabolism was a uniform and required component of malignancy (Warburg, 1956). This remained a major paradigm over the ensuing decades and was reinforced by studies in the 1980s-2000s demonstrating that oncogene expression (c-Myc, Ras, Akt, and others) is sufficient to stimulate glycolysis (Elstrom et al., 2004; Flier et al., 1987; Shim et al., 1997). It is now recognized that the tricarboxylic acid (TCA) cycle and other aspects of mitochondrial metabolism also contribute to cancer cell proliferation by providing biosynthetic precursors through the anaplerotic contributions of glucose, glutamine and other nutrients (Cheng et al., 2011; DeBerardinis et al., 2007; Weinberg et al., 2010). Other nutrients provide benefits to cancer cells through a variety of pathways, some of which are under oncogenic control (Boroughs and DeBerardinis, 2015). An emerging theme is that tumor cells acquire heterogeneous metabolic phenotypes to withstand complex challenges during cancer progression.

Cancer metabolism is regulated in part by changes in cell signaling and transcriptional programs activated by mutations in oncogenes and tumor suppressors, resulting in heterogeneous, cell-autonomous metabolic phenotypes among genetically diverse cancer cells (DeBerardinis and Chandel, 2016). Consistent with this idea, cultured cancer cells display variability in rates of nutrient uptake, with some activities associated with rapid cell proliferation (Jain et al., 2012). However, neither the full breadth of metabolic diversity in cancer cells, nor the complement of mechanisms by which tumor mutations elicit metabolic reprogramming, are known. We set out to characterize cell-autonomous metabolic heterogeneity in cancer cell lines derived from a particular tumor type. We chose non-small cell lung cancer (NSCLC) for the following reasons: *a)* NSCLC is the most prominent cause of cancer-related death worldwide, indicating that new therapeutics are needed; *b)* NSCLC cell lines covering the molecular diversity of this disease are easily available; *c)* many recurrently-mutated genes in NSCLC regulate metabolism, including *KRAS, EGFR, PIK3CA, TP53, KEAP1, PTEN* and others; and *d)* access to orthogonal high-content data sets from NSCLC cell lines would facilitate correlating metabolism with oncogenotypes, gene and protein expression and other features to understand the mechanisms by which metabolic phenotypes are established in these cells. A major goal was to understand whether these metabolic phenotypes predict vulnerability to targeted therapies or conventional chemotherapeutic agents.

## Results

### Experimental design and metabolic analyses

Over 80 highly-annotated NSCLC cell lines were analyzed for over 100 metabolic parameters derived from nutrient utilization, nutrient addiction, cell growth and isotope labeling after culture with [^13^C]glucose or [^13^C]glutamine (full dataset in **Supplemental Table S1**). The approach was designed to focus on glycolysis, glutaminolysis and the tricarboxylic acid (TCA) cycle, and to examine the relationship between these activities and orthogonal molecular and therapeutic sensitivity data (**Figure 1A**). In each experiment, multiple biological replicates cultured on different days were used to derive metabolic parameters. Consumption of glucose and glutamine and secretion of lactate and glutamate were measured to estimate rates of glycolysis and glutamine catabolism. Each cell line was assessed for proliferation in complete medium and for survival in media lacking either glucose or glutamine. Each cell line was also subjected to two complementary isotope labeling experiments designed to assess the extent to which glucose and glutamine supplied carbon to metabolite pools. One labeling experiment used medium with [U-^13^C]glucose and unlabeled glutamine, and the other used [U-^13^C]glutamine and unlabeled glucose. Metabolites were extracted after 6 and 24 hours and analyzed by gas chromatography–mass spectrometry (GC-MS) to obtain mass isotopologue distributions (MID) for citrate (Cit), fumarate (Fum), malate (Mal), lactate (Lac), serine (Ser) and glycine (Gly). Other than nutrient deprivation assays, the nutrient milieu was maintained across all experiments. These same conditions were used in experiments to characterize gene/protein expression and therapeutic sensitivity, making it possible to correlate our metabolic database with these other features.

**Figure 1.**
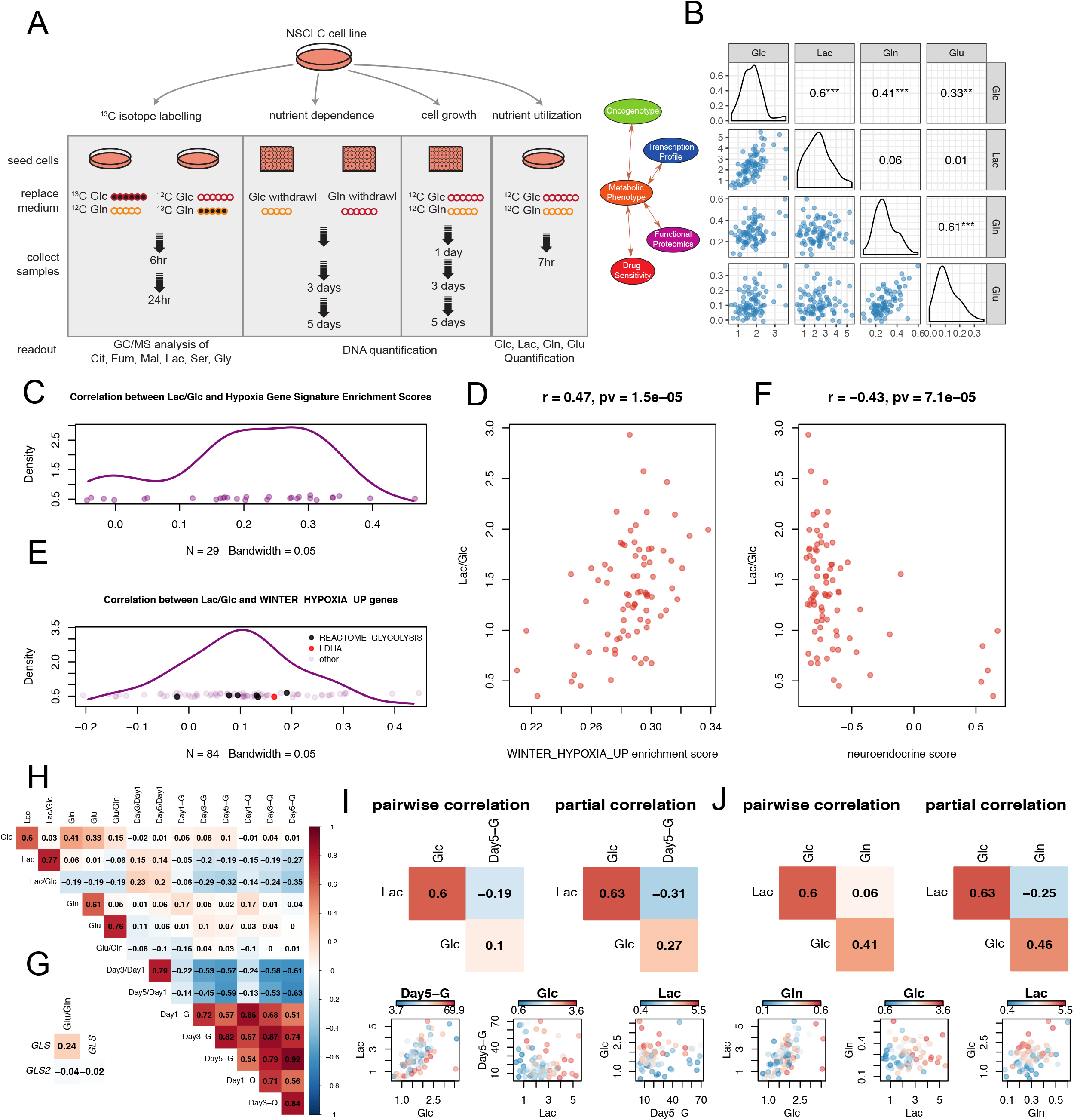
Experimental design and diversity of nutrient utilization in NSCLC cell lines. (**A**). Schematic diagram of experimental design for NSCLC metabolic profiling and downstream association analyses. Metabolic profiling data collected in our study include: isotopologue fractional enrichment in metabolites extracted from cells labeled with [U-^13^C]glucose or [U-^13^C]glutamine; cell survival on day 3 and 5 after withdrawal of glucose or glutamine; cell growth in complete medium on day 1, 3 and 5; glucose and glutamine consumption and lactate and glutamate secretion rates. Association analyses were performed to identify links between these metabolic features and orthogonal molecular phenotypes. See also Table S1. (**B**). Scatter plots and pairwise correlation amongst the nutrient utilization features. Correlation coefficients were derived from Pearson correlations. ***, p<=0.001; **, p<=0.01; *, p<=0.05; no asterisk is used when p>0.05. (**C**). Kernel density estimation of correlation coefficient distribution from the pairwise correlations between Lac/Glc ratio and ssGSEA scores from 29 hypoxia-related genesets. The vast majority (25/29) of the correlations are positive. The hypoxia related genesets were selected from C2-CGP gene sets in MSigDB with the criterion that the geneset name contained “HYPOXIA” but not “DN” (short for “Down”). (**D**). Scatterplot showing the positive correlation between Lac/Glc and ssGSEA scores derived for “WINTER_HYPOXIA_UP” signature. (**E**). Kernel density estimation of correlation coefficient distribution from the pairwise correlation between Lac/Glc ratio and 84 genes from the “WINTER_HYPOXIA_UP” gene set. Glycolytic genes (*ALDOA, PGK1, TPI1, PGAM1, GAPDH* and *PFKFB4*) from “REACTOME_GLYCOLYSIS” geneset and *LDHA* are indicated by black and red spots. (**F**). Scatterplot showing the negative correlation between Lac/Glc and neuroendocrine scores derived from NSCLC cell lines. (**G**). Pearson correlation between glutaminolysis (Glu/Gln) and expression of *GLS* and *GLS2*. A positive correlation is only observed between Glu/Gln and *GLS* expression. (**H**). Correlation heatmap revealing pairwise Pearson correlations amongst metabolic features from nutrient utilization, cell growth and nutrient dependency. (**I** and **J**). Comparisons of partial correlations and pairwise correlations for Lac, Glc and Day5-G (**I**) or Lac, Glc and Gln (**J**). Abbreviations: Lac, lactate; Glc, glucose; Gln, glutamine; Glu, glutamate.

### Nutrient consumption, secretion and dependence

Nutrient consumption/secretion rates were variable across the cell lines, varying approximately 6-7-fold for glucose consumption and 15-fold for lactate secretion (**Supplementary Table 1**). As expected, glucose consumption correlated strongly with lactate secretion, because lactate is the major metabolic product of glucose in culture; similarly, glutamine consumption correlated strongly with glutamate secretion (**Figure 1B**). On average, cells secreted 1.35 moles of lactate per mole of glucose consumed (standard deviation 0.5 moles), and 0.4 moles of glutamate per mole of glutamine consumed (standard deviation 0.2 moles), emphasizing that cultured cancer cells process both glucose and glutamine at rates far exceeding their need to retain carbon from these nutrients (**Supplementary Table 1**). A positive correlation was also observed between glucose and glutamine consumption, indicating an unexpected coordination between glutaminolysis and glucose consumption (**Figure 1B**).

Normalizing lactate secretion to glucose consumption (Lac/Glc ratio) provides an estimate of each cell line’s preference for aerobic glycolysis. The cell lines displayed an 8-fold range of Lac/Glc ratios, indicating that NSCLC cell lines are diverse in the extent to which they discard glucose carbon as lactate (**Supplementary Table 1**). The Lac/Glc ratio correlated significantly but weakly with a measure of cell proliferation in complete medium (Day3/Day1 ratio; p=0.04) (**Figure 1H**). To identify associations between Lac/Glc and transcriptional programs, we derived single sample gene set enrichment analysis (ssGSEA) scores for each cell line based on the C2-CGP gene sets representing expression signatures of genetic and chemical perturbations from MSigDB (Subramanian et al., 2005). Enrichment scores from ssGSEA represent the degree to which genes in a particular gene set are coordinately up- or down-regulated (Barbie et al., 2009). We found that the vast majority (25/29) of enrichment scores previously generated by hypoxia-related signatures are positively associated with the Lac/Glc feature (**Figure 1C**); an example gene set is shown in **Figure 1D** (Winter et al., 2007). This gene set includes *LDHA* and several other glycolytic genes as defined by REACTOME (Fabregat et al., 2018; van Wijk and van Solinge, 2005). Individual members of this gene set exhibit a moderate but overall positive correlation with Lac/Glc (**Figure 1E**). On the other hand, a negative correlation was observed between Lac/Glc and ssGSEA scores derived from gene sets related to neuronal processes, which tend to be expressed in cell lines with neuroendocrine differentiation (Ionescu et al., 2007). We therefore derived neuroendocrine scores for our cell lines based on a 50-gene signature (Zhang et al., 2018) and found that cell lines with high neuroendocrine scores are invariably low in Lac/Glc (**Figure 1F**). Altogether these data indicate that hypoxia gene sets correlate with glycolytic metabolism even when cells are cultured in normoxia, and that cell lines with neuroendocrine-like gene expression signatures release relatively little lactate per glucose consumed.

We used a similar approach to normalize glutamate secretion to glutamine uptake (Glu/Gln ratio) as a surrogate for the release of carbon derived from glutamine. Glu/Gln was also heterogeneous among the cell lines, ranging from essentially no glutamate released to a 1:1 molar ratio of glutamate release per glutamine consumed. Glu/Gln correlates positively with *GLS* mRNA, which encodes the key glutaminolytic enzyme GLS (**Figure 1G**). *GLS2*, which encodes another glutaminase isoform, did not correlate with Glu/Gln at the transcript level, and *GLS* did not correlate with *GLS2*.

To understand relationships between nutrient utilization, nutrient dependency and cell growth, we examined pairwise correlations amongst these features in a correlation heatmap (**Figure 1H**). The two cell growth features (Day3/Day1 and Day5/Day1) are strongly correlated, as expected. The six features related to glucose and glutamine dependence (Day1-G, Day3-G, Day5-G, Day1-Q, Day3-Q, Day5-Q) also correlate with each other, suggesting that cells sensitive to glucose deprivation also tend to be sensitive to glutamine deprivation, and vice versa. Growth rate correlates negatively with nutrient dependence, indicating that cell lines which grow rapidly under nutrient replete conditions perish more rapidly upon either glucose or glutamine withdrawal. However, neither glucose nor glutamine consumption correlate with cell growth rates, suggesting that these nutrients are used for processes other than or in addition to biomass assimilation.

In some cases, partial correlation produced stronger associations among metabolic features than direct pairwise correlations. **Figure 1I** provides an example where the negative association between Lac and Day5-G is more significant in a partial correlation controlling for Glc than in a pairwise correlation, indicating that for a given rate of glucose uptake, the more lactate produced, the more sensitive the cell line is to glucose deprivation. In **Figure 1J**, the negative association between Lac and Gln is more significant in a partial correlation controlling for Glc than in a pairwise correlation. In other words, with the same amount of glucose uptake, the more lactate a cell produces, the less glutamine it consumes.

### Diversity in metabolic pathway utilization inferred by mass isotopologue distributions

Mass isotopologue distributions (MIDs) report the metabolic fates of ^13^C-labeled fuels, providing a view of metabolism that cannot be achieved from steady-state metabolite levels (Buescher et al., 2015; Jang et al., 2018). MID analysis has been used extensively to assess various forms of the TCA cycle involving differential nutrient contributions to metabolic intermediates in this pathway. These pathways are controlled by both cell-intrinsic and cell-extrinsic influences. For example, oxidative and reductive pathways of glutamine metabolism are readily differentiated using MID of citrate following culture with [U-^13^C]glutamine. Glutamine oxidation, a major source of anaplerosis in cancer cell lines, generates m+5 labeling in α-ketoglutarate and m+4 labeling in other TCA cycle metabolites (**Figure 2A**). Glutamine-dependent reductive carboxylation (GDRC) generates α-ketoglutarate m+5, citrate m+5, acetyl-CoA m+2, and m+3 labeling in other TCA cycle intermediates (**Figure 2B**). Several processes relevant to tumor biology impair glutamine oxidation and enhance labeling from GDRC; these include hypoxia, *VHL* mutation and/or suppression of various components of oxidative metabolism through mutation or pharmacological inhibition (Gameiro et al., 2013; Metallo et al., 2011; Mullen et al., 2011; Rajagopalan et al., 2015).

**Figure 2.**
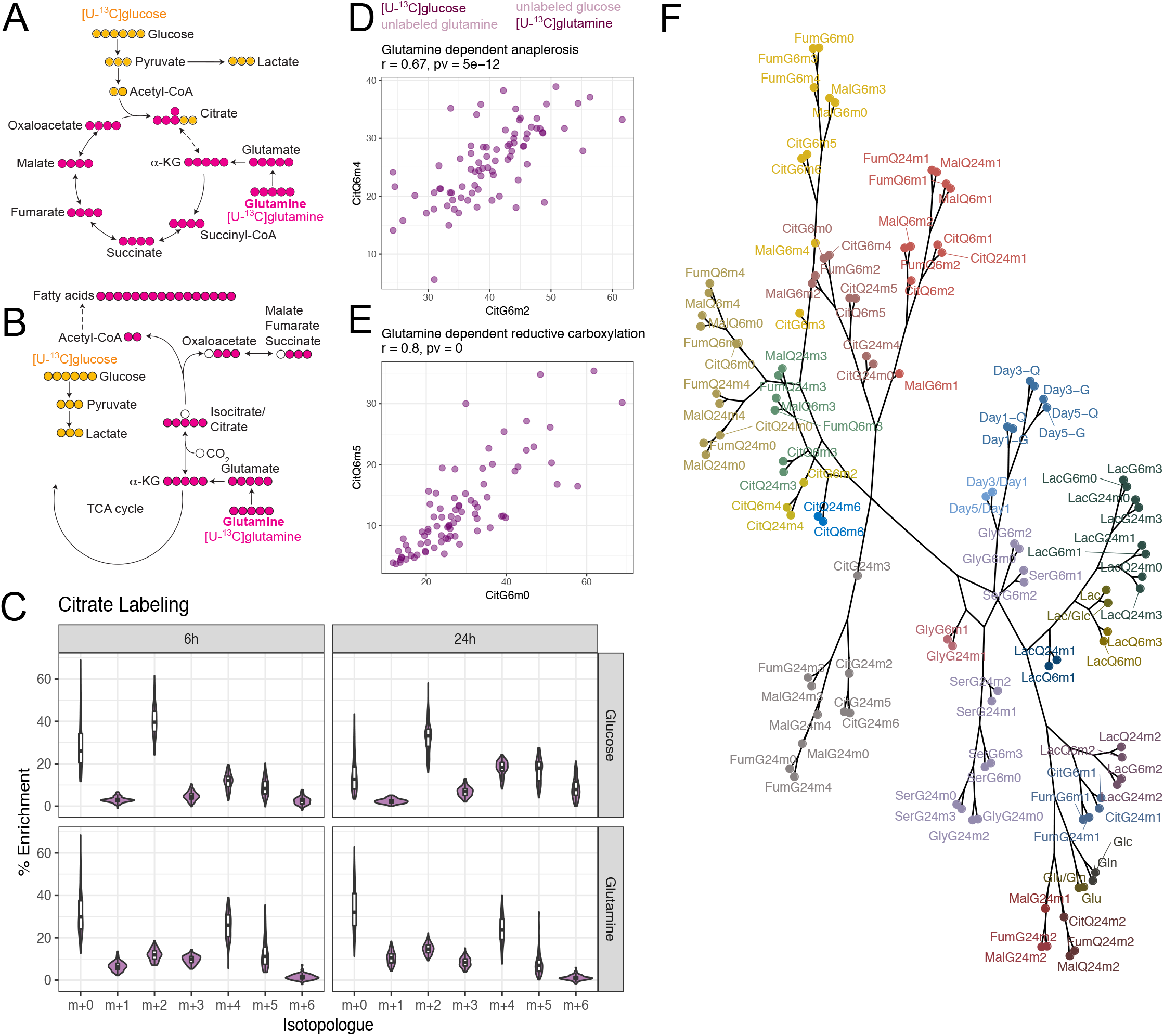
Differential pathway utilization inferred by mass isotopologue distributions. (**A** and **B**). Schematic diagrams of representative isotopologues generated from [U-^13^C]glucose or [U-^13^C]glutamine labeling through glutamine-dependent anaplerosis (**A**) or glutamine-dependent reductive carboxylation (**B**). (**C**). Violin plots showing distribution of citrate isotopologues from different tracers and different labeling durations (6h and 24h). See also Figure S2. (**D** and **E**). Scatterplots showing that enrichment of signature isotopologues produced from two different tracers for the same metabolic pathway (glutamine-dependent anaplerosis in **D** and glutamine-dependent reductive carboxylation in **E**) are highly correlated with each other. (**F**). Dendrogram of all metabolic features from hierarchical clustering with absolute Pearson correlation-based distance using Ward’s minimum variance method. The 23 branches were arbitrarily colored. The dendrogram follows a phylogenic tree layout to allow easy identification of closely related metabolic features. Abbreviations: Cit, citrate; Fum, fumarate; Mal, malate; Lac, lactate; Ser, serine; Gly, glycine.

To our knowledge, there have been no previous attempts to use parallel-tracer MID analysis to capture metabolic diversity in a large panel of cell lines derived from one type of cancer. We named ^13^C labeling features using an abbreviation for the metabolite, followed by the tracer, labeling duration and number of ^13^C nuclei. For example, m+2 citrate after 6 hours of [U-^13^C]glucose labeling is named CitG6m2. An initial examination of the data revealed remarkable cell-autonomous diversity in isotope labeling among the cell lines, but good consistency among biological replicates (**Figure S1**).

Examination of MIDs in citrate after labeling with either [U-^13^C]glucose or [U-^13^C]glutamine reveals several interesting features (**Figure 2C**). First, the distribution of MIDs is generally conserved between the two time points, indicating that most labeling occurs within the first 6 hours. The modest changes at 24 hours included reduced prominence of unlabeled citrate and increased abundance of isotopologues derived from multiple rounds of the TCA cycle (e.g. m+4 and m+6 from [U-^13^C]glucose). Second, labeling patterns are heterogeneous across the panel, with some citrate MIDs (particularly those involving m+0 and m+2 from [U-^13^C]glucose and m+0, m+4 and m+5 from [U-^13^C]glutamine) covering about one third of the total possible MID range. This diversity indicates heterogeneity of pathway preference in these cell lines, even when cultured under identical conditions.

Third, the abundant m2 isotopologues from [U-^13^C]glucose at 6 and 24 hours, coupled with the m4 isotopologues from [U-^13^C]glutamine, demonstrate the expected prominence of glucose-dependent acetyl-CoA formation and glutamine-dependent anaplerosis (DeBerardinis et al., 2007). In this common pathway, glucose and glutamine metabolism converge on citrate synthesis (**Figure 2A**). Although both CitGlc6m2 and CitQ6m4 fractions varied widely across the panel, they correlated well with each other (**Figure 2D**). MIDs from other TCA cycle intermediates were also diverse and agreed with the prominent pattern of glutamine-dependent anaplerosis across the panel (**Figure S2A-E;** standard deviations are in **Figure S2F**). Other pairs of glucose- and glutamine-derived isotopologues also demonstrated strong correlations. One of the most striking was the positive correlation between CitQ6m5 and CitGlc6m0, two isotopologues characteristic of GDRC (**Figure 2E**). Surprisingly, some cells had as much as 35% m5 labeling in citrate from [U-^13^C]glutamine; this was unexpected considering that GDRC has largely been observed in hypoxia and other conditions of impaired glucose and glutamine oxidation.

Relationships among MIDs at 6h and other metabolic features are displayed as a correlation heatmap in **Figure S3**, with the results summarized in **Tables S2.1** and **S2.2**. We also generated a dendrogram from hierarchical clustering using absolute correlation-based distances for each feature (**Figure 2F**). In general, the same isotopologues are strongly correlated at 6h and at 24h (**Figure S3, Tables S2.1** and **S2.2**). Stronger correlations are observed for isotopologues with a larger dynamic range. Strong correlations are also enriched between isotopologues from metabolites in the same metabolic pathway. Examples include shared labeling features among TCA cycle intermediates (citrate, fumarate and malate) or between serine and glycine. Surprisingly, there was a paucity of associations between ^13^C labeling and features related to nutrient utilization, cell proliferation and nutrient dependence. This indicates that even though many pathways reported by ^13^C labeling contribute to cell proliferation, the prominence of a given set of labeling features predicts neither the growth rate nor nutrient dependence. Despite the overall paucity of associations between ^13^C and non-^13^C features, we did capture some expected associations, including a positive correlation between Lac and LacGm3 (**Figure S3**).

### Glucose-dependent anaplerosis inferred from [U-^13^C]glucose labeling predicts dependence on pyruvate carboxylase

Alternative positional labeling of precursors can simplify interpretation of labeling in metabolic products. For example, [3,4-^13^C]glucose is a preferred tracer to measure glucose-dependent anaplerosis via pyruvate carboxylase (PC). This tracer differentiates PC-dependent vs. PDH-dependent entry of glucose carbon into the TCA cycle, because [3,4-^13^C]glucose is converted to [1-^13^C]pyruvate. PDH removes the labeled carbon as ^13^CO_2_, whereas PC transfers the label to OAA so that it is retained in TCA cycle intermediates as m+1 isotopologues (**Figure 3A**). When [U-^13^C]glucose is the precursor, glycolysis produces [U-^13^C]pyruvate, and both PDH and PC transfer label from [U-^13^C]pyruvate to the TCA cycle, producing m+2 or m+3 isotopologues on the first turn. However, combined activity of PDH and PC, coupled with multiple turns of the cycle, complicate interpretation of MIDs from [U-^13^C]glucose (**Figure 3B** and **S4**). It would be advantageous to derive information about PC from [U-^13^C]glucose, because *a)* anaplerosis is an important component of biomass assimilation and cell growth; *b)* PC is the primary anaplerotic route in some tumors in vivo and accounts for glutamine-independent growth in some cell lines (Cheng et al., 2011; Sellers et al., 2015); and c) the high cost of [3,4-^13^C]glucose relative to [U-^13^C]glucose makes the former tracer challenging to use for human in vivo experiments.

**Figure 3.**
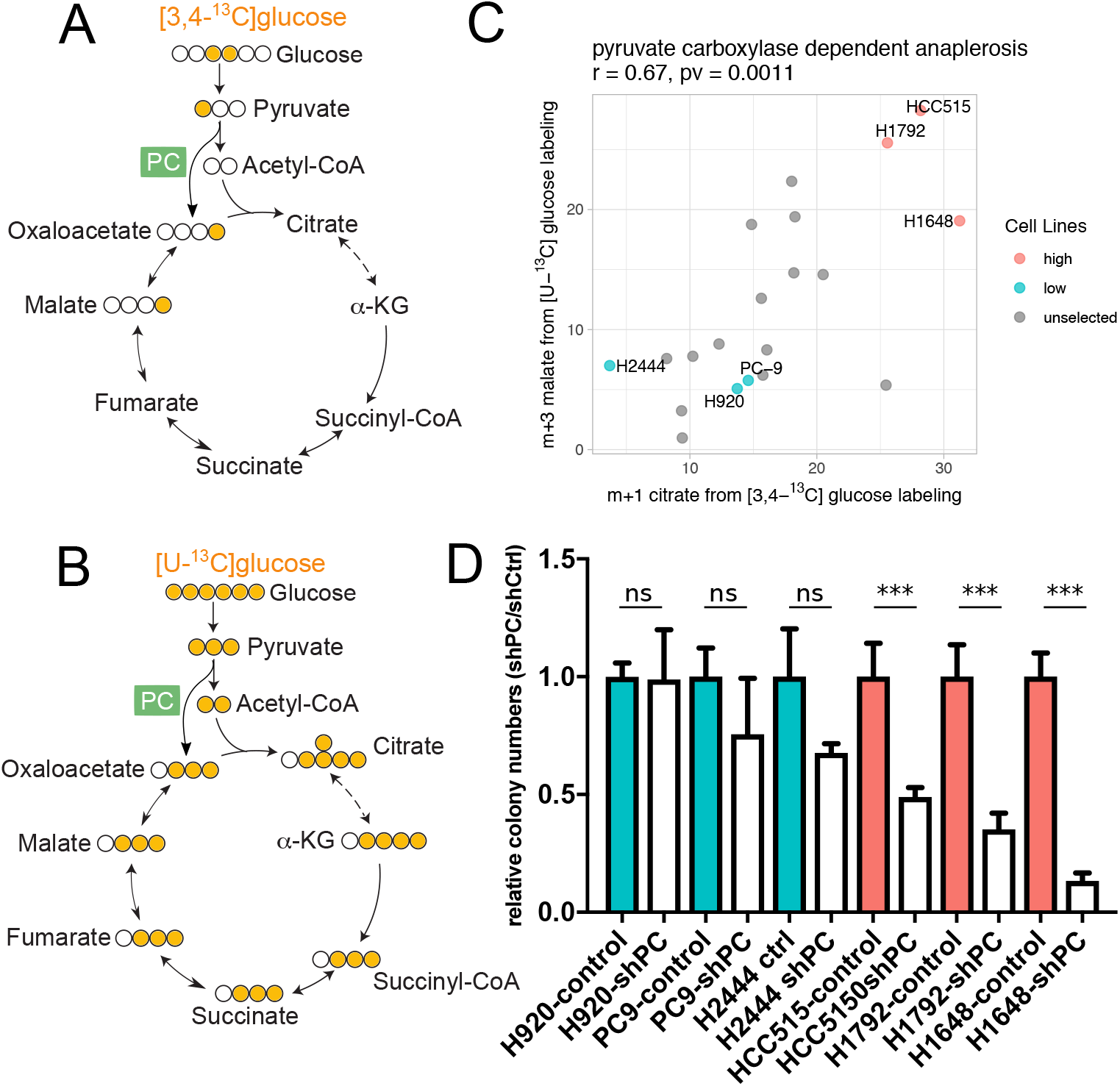
Inferring pyruvate carboxylase activity from enrichment of malate isotopologues. (**A** and **B**). Signature isotopologues produced from pyruvate carboxylation with [3,4-^13^C]glucose (**A**) or [U-^13^C]glucose (**B**) labeling. (**C**). Scatter plot showing positive correlation between m+1 citrate from [3,4-^13^C]glucose labeling and m+3 malate from [U-^13^C]glucose labeling. Both these isotopologues are presumably generated through pyruvate carboxylation. Cell lines selected for PC dependency testing in (**D**) are indicated with labels. Correlation coefficient and p-value from Pearson correlation are provided in title. (**D**). Effect of PC silencing on colony formation in cells lines predicted to have high (red) or low (blue) PC-dependent anaplerosis based on the data in (**C**). Data are represented as mean ± SEM. Statistical significance based on t-test, ***p < 0.005. See also Figure S4.

To test whether [U-^13^C]glucose can reliably report PC activity, we cultured a subset of 20 cell lines in [3,4-^13^C]glucose and compared the resulting m+1 isotopogues to m+3 isotopologues derived from [U-^13^C]glucose. Specifically, we compared citrate m+1 with malate m+3; the latter species arises through malate dehydrogenase-dependent exchanges with OAA m+3, which was not abundant enough to detect in our samples. The strong positive correlation between these isotopologues indicates that [U-^13^C]glucose reports PC activity (**Figure 3C**). MalG6m3 also correlates with the abundance of PC mRNA (**Figure S4C**). To remove the confounding effect of MalG6m3 arising from multiple turns of the TCA cycle, we fitted a linear regression model using CitG6m4, which also arises from multiple TCA cycle turns, to predict the portion of MalG6m3 arising independently of PC. The residuals from this fit, which we take to arise from PC activity, correlate much better with *PC* mRNA expression (**Figure S4D**).

To assess whether cancer cells with higher PC activity also had enhanced dependence on the enzyme, we selected three cell lines each with high (HCC515, H1792, H1648) or low (H920, PC9, H2444) PC activity as reported by labeling from [U-^13^C]glucose. Each cell line was modified to express a control shRNA or an shRNA directed against PC. PC silencing reduced soft agar colony formation in cells with high PC-dependent labeling, but not in cell lines with low PC-dependent labeling (**Figure 3D**). Together, these data indicate that mass isotopologues from [U-^13^C]glucose can be used to predict PC activity, gene expression and dependence.

### Associations between MIDs and oncogenotypes

Next we explored relationships between metabolic phenotypes and NSCLC oncogenotypes. We clustered cell lines based on citrate MIDs after [U-^13^C]glucose labeling for 6h and examined mutations in recurrently-mutated NSCLC genes thought to regulate metabolism, including *EGFR, KRAS* and *STK11* (**Figure 4A**). Generally, stronger associations were observed when site-specific rather than site-agnostic mutations were considered (**Figure 4A**). Examination of unlabeled citrate (CitGlc6m0), the isotopologue with the largest dynamic range in this dataset, demonstrates this point. Cells with *EGFR* mutations tended to have higher fractional contents of CitGlc6m0, but this relationship was particularly striking for exon 19 deletions (**Figure 4A,B**). To test whether this association translates to primary human NSCLC, we used our database of intra-operative [U-^13^C]glucose infusions in NSCLC patients. In these clinical studies, patients with solitary pulmonary lesions are subjected to [U-^13^C]glucose infusion during surgical resection of the tumor and adjacent lung (Faubert et al., 2017; Hensley et al., 2016). Fragments from six *EGFR*-mutant tumors had significantly higher Citm0 fractions than tumors with wild-type *EGFR* (**Figure 4F**). In contrast, no differences in labeling were noted for adjacent lung samples from the same patients (**Figure 4F**). These findings indicate that some subtype-selective labeling phenotypes are observed both in culture and in primary human tumors.

**Figure 4.**
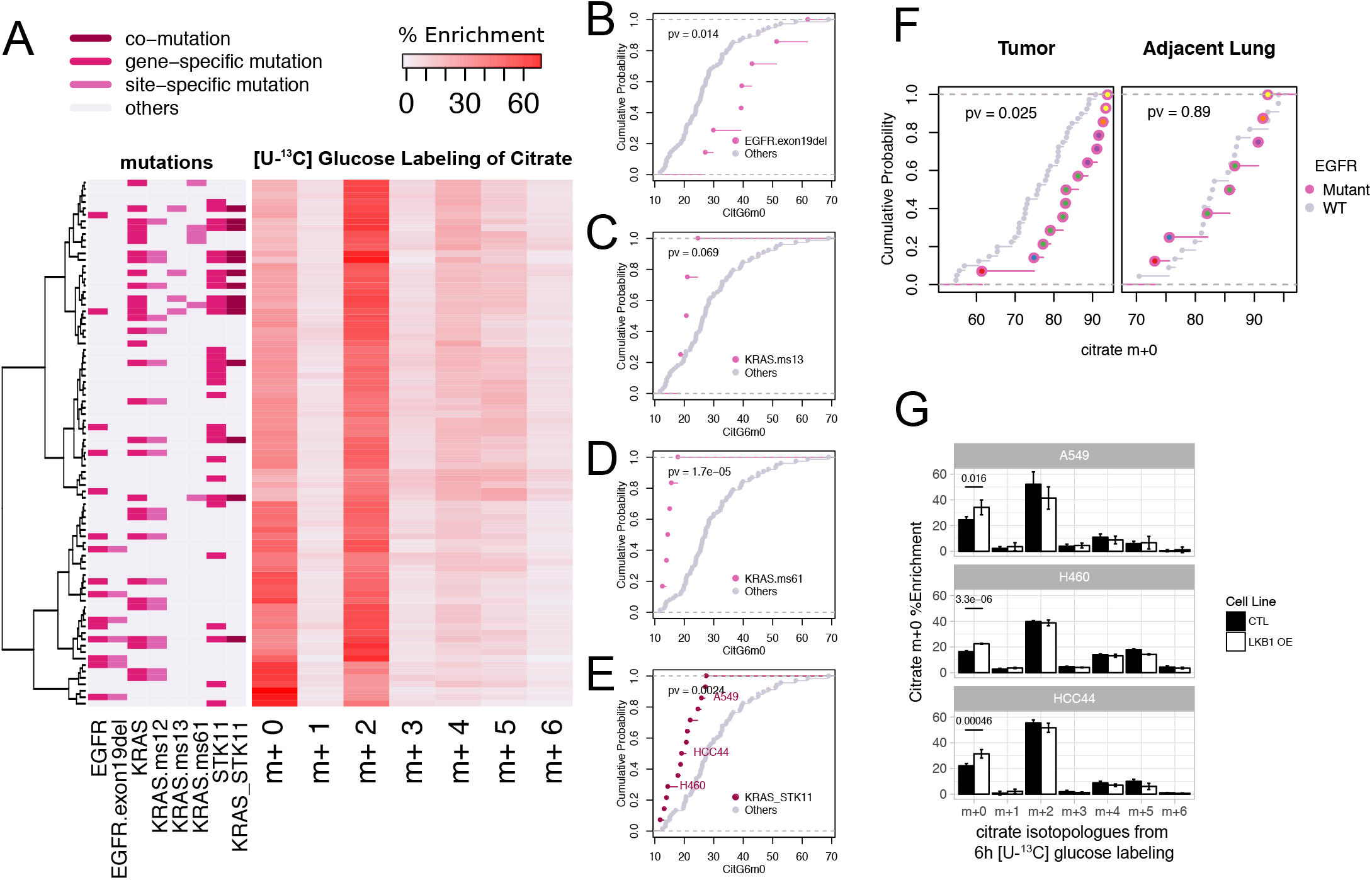
Associations between oncogenotypes and citrate mass isotopologues. (**A**). Heatmap with hierarchical clustering of cell lines and citrate mass isotopologues from [U-^13^C]glucose labeling for 6h. Clustering was based on Ward’s minimum variance method. Relevant oncogenotypes are indicated. (**B-E**). Cumulative distribution function plots showing different levels of CitG6m0 in cell lines with specific oncogenotypes (*EGFR* exon 19 deletion in (**B**); *KRAS* missense mutation of the 13^th^ codon in (**C**); *KRAS* missense mutation of the 61^st^ codon in (**D**); Concurrent *KRAS* and *STK11* mutation in (**E**). compared to the rest of the cell lines (p-values are generated based on Kolmogorov–Smirnov test). (**F**). *Left*, cumulative distribution function plots comparing citrate m+0 fractions between tumor fragments with *EGFR* mutations and tumor fragments lacking *EGFR* mutations. *Right*, cumulative distribution function plots of the adjacent lung samples from these same patients. Note that for the tumors we compared 14 fragments from 6 patients with *EGFR* mutations to 40 fragments from 22 *EGFR* WT patients, whereas for the adjacent lung tissues, we compared 8 fragments from 6 patients with *EGFR* mutations to 22 fragments from 22 *EGFR* WT patients. Patient origins of the *EGFR* mutant fragments are indicated by different colors inside the circles. (**G**). Re-expression of LKB1 (encoded by *STK11*) in three selected cell lines with co-mutant *KRAS* and *STK11* increases the citrate m+0 fraction after labeling with [U-^13^C]glucose for 6h (**G**). Data are represented as mean ± SEM. See also Figure S5. OE, over-expression.

Cell lines with KRAS missense mutations at Q61 have low CitGlc6m0, and KRAS G13 mutants display a similar trend (**Figure 4A,C,D**). *KRAS-mutant* cell lines with concurrent loss of function *STK11* mutations have distinct metabolic vulnerabilities from cell lines with mutations in *KRAS* alone (Kim et al., 2017; Liu et al., 2013), and tumors with this combination of mutations have enhanced aggressiveness in mice and humans (Calles et al., 2015; Ji et al., 2007). We find that cell lines with concurrent *KRAS/STK11* mutations have low CitGlc6m0 content, indicating a propensity to supply the TCA cycle with glucose carbon (**Figure 4E**). Re-constituting each of three independent co-mutant cell lines with wild-type *STK11* resulted in a small but significant increase in the CitGlc6m0 fraction, indicating that *STK11* regulates glucose’s contribution to the TCA cycle under these conditions (**Figure 4G, S5**). This observation is consistent with previous reports focusing on *STK11’s* role in regulating nutrient metabolism (Faubert et al., 2014; Kim et al., 2013).

### GDRC is associated with an epithelial state and sensitivity to EGFR inhibitors

The unexpected prominence of GDRC labeling in some NSCLC cells (**Figure 2E**) prompted us to examine orthogonal data for novel associations with this pathway. We used GSEA to identify transcriptional programs associated with GDRC, focusing on CitG6m0 and CitQ6m5. This analysis yielded strong positive associations with an epithelial gene set targeted by the epithelial-mesenchymal transition (EMT)-related transcription repressor ZEB1 (Aigner et al., 2007) and negative associations with a gene set related to resistance to Gefitinib, an EGFR inhibitor (Coldren et al., 2006) (**Figure 5A,B** and **Figure S6A,B**). These associations implied a relationship between GDRC and an epithelial phenotype characterized by EGFR signaling. Consistent with this idea, data from reversed-phase proteomics arrays (RPPA) revealed positive correlations between GDRC and both β-catenin and E-cadherin, two components of the catenin-cadherin complex critical for maintaining epithelial integrity (**Figure 5C**). GDRC also correlated positively with EGFR phosphorylation at Y1173 (pY1173), a marker for EGFR signaling and Gefitinib sensitivity (**Figure 5C**). Finally, comparing metabolic features with high-throughput drug sensitivity data (McMillan et al., 2018) revealed that GDRC cell lines tend to be sensitive to the EGFR inhibitor Erlotinib; these cell lines have low areas under the Erlotinib dose-response curve (AUC) (**Figure 5C**).

**Figure 5.**
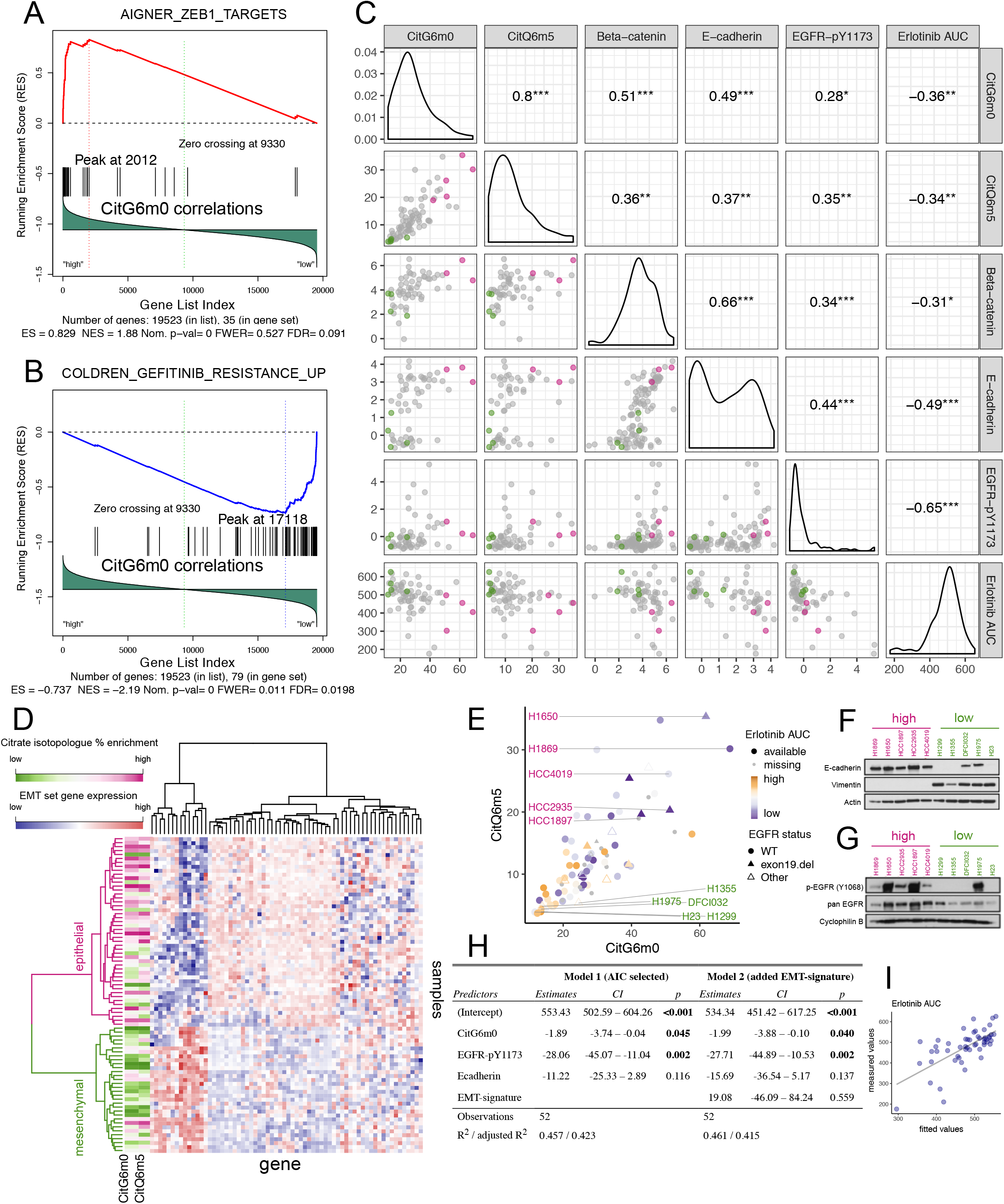
Reductive carboxylation is associated with an epithelial state and is enriched in cell lines sensitive to EGFR inhibitors. (**A-B**). Gene Set Enrichment Analysis (GSEA) identified CitG6m0 as positively correlated with ZEB1 target genes (**A**) and negatively correlated with Gefitinib resistance genes (**B**). (**C**). Scatterplot and pairwise Pearson correlation among GDRC metabolic features CitG6m0, CitQ6m5; RPPA features beta-catenin, E-cadherin and EGFR-pY1173; and compound sensitivity feature Erlotinib AUC (higher area under the dosing curve represents higher resistance). The color scheme for points in the scatterplot is explained in the legend for (**E**). *** denotes p<=0.001; ** denotes p<=0.01; * denotes p<=0.05; no asterisk is used when p>0.05. (**D**). Heatmap with hierarchical clustering of samples and EMT signature genes. Clustering was based on Ward’s minimum variance method. The CitG6m0 and CitQ6m5 fractions are indicated by the color scale. Higher levels of GDRC metabolic features were observed for the epithelial cluster. (**E**). Scatter plot of CitG6m0 and CitQ6m5 with *EGFR* mutation status marked by different symbols and EGFR inhibitor sensitivity indicated by different color and shapes. Five GDRC-high cell lines and five GDRC-low cell lines were selected for further characterization. These cell lines are also indicated by coloring in (**C**). (**F-G**). Validation of EMT status and EGFR activation by western blotting for the 10 selected cell lines. In (**F**), the epithelial marker E-cadherin is expressed in GDRC-high cell lines, whereas the mesenchymal marker vimentin is expressed GDRC-low cell lines. In (**G**), p-EGFR (Y1068) indicative of EGFR activation is more prominent in GDRC-high cell lines. (**H**). Coefficients and p-values from multiple regression models predicting inhibitor sensitivity from different feature sets. Model 1 was obtained from stepwise feature selection based on Akaike information criterion (AIC) with input features including CitG6m0, CitQ6m5, EGFR-pY1173, E-cadherin, beta-catenin and EMT class. Model 2 adds the EMT-signature (EMT class) into Model 1. Note that the p-values for CitG6m0 are significant in both models while controlling for the RPPA features or gene expression-derived EMT feature. (**I**). Scatterplot of fitted values from model 1 in (**H**) and the measured value (Erlotinib AUC). See also Figure S6.

Sensitivity to EGFR inhibitors is predicted by an EMT signature defined by expression of 76 genes, with mesenchymal cells demonstrating resistance to EGFR inhibitors regardless of their *EGFR* mutation status (Byers et al., 2013). Using the reported EMT signature to cluster our cells into mesenchymal-like and epithelial-like groups, we found that cell lines with high levels of GDRC-related isotopologues were overrepresented in the epithelial-like group (**Figure 5D** and **Figure S6C**). Lac/Glc and Glu/Gln features were similar between the two groups. We selected five GDRC-high and five GDRC-low cell lines differing by *EGFR* mutation status for further analysis (**Figure 5E** and **Table S3**). On western blotting, GDRC-high cell lines had high E-cadherin and low vimentin levels, consistent with the predicted epithelial-like state (**Figure 5F**). They also had abundant EGFR Y1068 phosphorylation, despite the fact that only two (H1650 and HCC1935) had detectable *EGFR* mutations (**Figure 5G**). In the GDRC-low lines, Y1068 phosphorylation was only observed in H1975, one of the two *EGFR*-mutant cell lines we selected in this class (**Figure 5G**). We also assessed the correlation between CitGlc6m0 and E-cadherin at the level of gene methylation, gene expression and protein expression, and found the best correlation with protein expression (**Figure S6D**).

To exploit the potential of metabolic features as predictors of EGFR inhibitor sensitivity, we fitted a multiple regression model starting from a feature set including GDRC-related isotopologues (CitG6m0 and CitQ6m5); the binary epithelial/mesenchymal classification based on EMT signature clustering; and the RPPA features EGFR-pY1173, E-cadherin and beta-catenin. Then we performed stepwise feature selection, with the model selected using Akaike information criterion (AIC). Features remaining in the final multiple regression model included CitG6m0, EGFR-pY1173 and E-cadherin, with significant p-values for CitG6m0 and EGFR-pY1173 (**Figure 5H**, Model 1 and **Figure 5I**). Importantly, when the EMT-signature binary classification feature was added to the model, the p-value for CitG6m0 remained significant, indicating that this metabolic feature predicts EGFR inhibitor sensitivity independently of features based on gene and protein expression (**Figure 5H**, Model2).

### *De novo* serine synthesis is associated with pemetrexed sensitivity

Serine biosynthesis has garnered increasing attention because of serine’s multifaceted roles in methionine, lipid, folate and nucleotide metabolism, as well as in amino acid transport, all of which are important in cancer cells (Mattaini et al., 2016). During *de novo* serine synthesis, [U-^13^C]glucose is converted to serine m+3 via the glycolytic intermediate 3-phosphoglycerate (3-PG). Serine m+3 is then converted to glycine m+2 by serine hydroxymethyltransferases (SHMT), resulting in a high correlation between serine SerG6m3 and GlyG6m2 isotopologues from our ^13^C tracing data (**Figure 6A**). A subset of these labeling data were previously used to discover regulation of serine biosynthesis by NRF2 and ATF4 (DeNicola et al., 2015). Indeed, many transcripts involved in serine biosynthesis and related pathways are highly correlated with the SerG6m3 isotopologue (**Figure S7A**).

**Figure 6.**
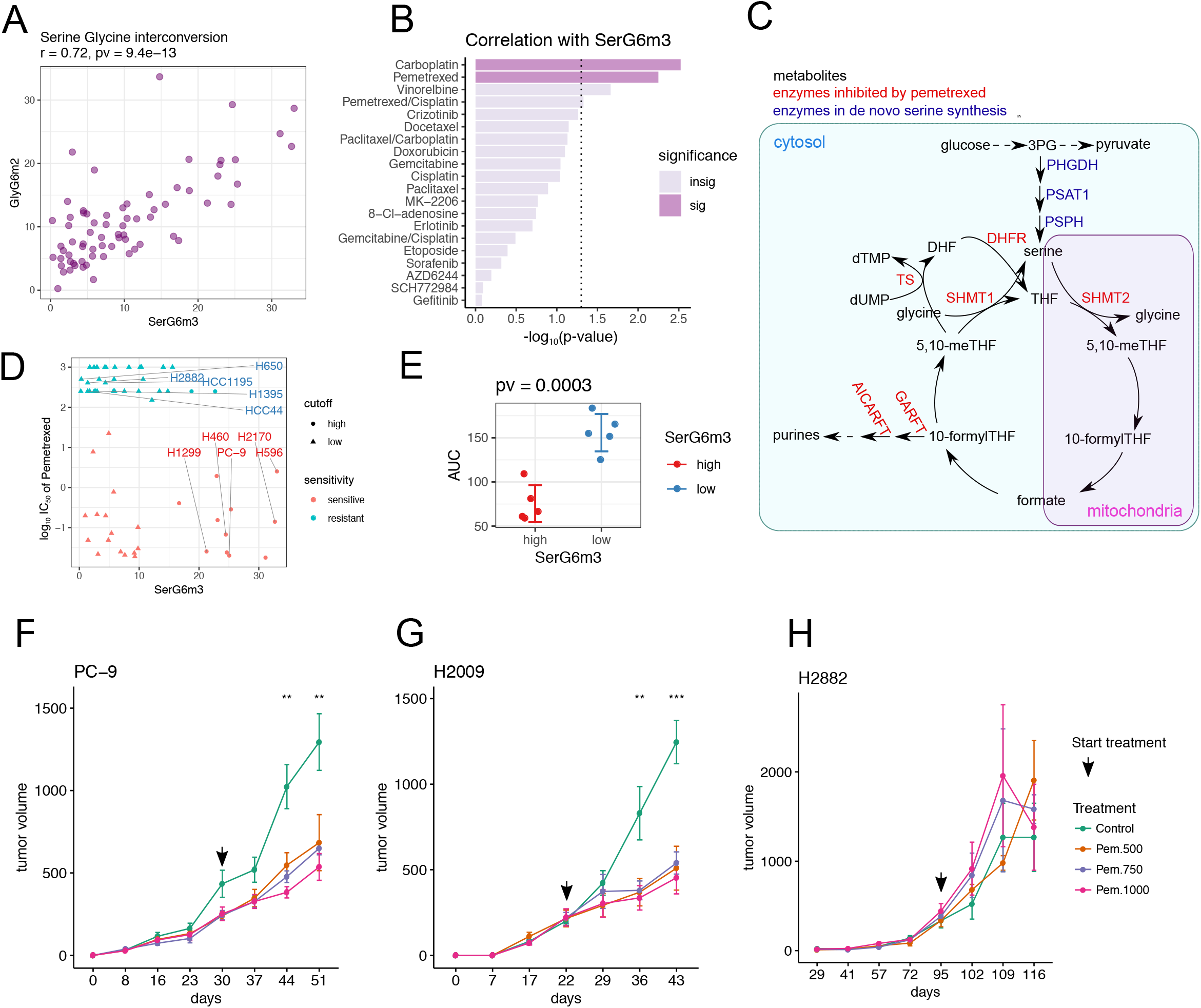
*De novo* serine synthesis from glucose is associated with sensitivity to pemetrexed. (**A**). Scatterplot showing positive correlation between SerG6m3 and GlyG6m2. Correlation coefficient and p-value from Pearson correlation are provided in the title. (**B**). Drug sensitivity correlations with SerG6m3. −log10 (p-values) from Pearson correlations are plotted and the hits were ranked by decreasing statistical significance. The dashed line demarcates the nominal p-value cut-off of 0.05 after −log10 transformation, and the darkly-colored bars denote statistical significance after multiple comparison adjustment by Benjamin Hochberg procedures. (**C**). Schematic diagram of the serine biosynthesis pathway feeding into one-carbon metabolism. Metabolites are in black, serine *de novo* synthesis enzymes are in blue and enzymes reportedly targeted by pemetrexed are in red. (**D**). Relationship between SerG6m3 and pemetrexed IC50, and selection of cell lines for further characterization. Note that almost all the cell lines with high SerG6m3 are sensitive to pemetrexed. Cell lines are colored based on pemetrexed sensitivity. 5 pemetrexed sensitive cell lines with high SerG6m3 and 5 pemetrexed resistant cell lines with low SerG6m3 were selected for further characterization. (**E**). Validation of pemetrexed sensitivity between selected cell lines with high and low SerG6m3 fractions. P-value from t-test is displayed in the title. See also Figure S7. (**F-H**). In vivo testing of pemetrexed sensitivity in xenograft mouse models. PC-9 (**F**) and H2009 (**G**), cell lines sensitive to pemetrexed in culture, are also sensitive to pemetrexed in vivo, whereas the pemetrexed-resistant cell line H2882 (**I**) retains this resistance in vivo. The arrow indicates the time when pemetrexed therapy was initiated. Data are represented as mean ± SEM. Statistical significance at each time point was determined by one-way ANOVA.

A search for correlations between SerG6m3 and therapeutic sensitivities (Table S4) identified the antifolate pemetrexed as one of the top hits among many that targets cell replication (**Figure 6B**). This association was interesting because pemetrexed inhibits several folate-dependent reactions from one-carbon metabolism and hence impacts nucleotide biosynthesis (Curtin and Hughes, 2001; Daidone et al., 2011; Ducker and Rabinowitz, 2017) (**Figure 6C**). We selected 5 pemetrexed-sensitive/SerG6m3-high cell lines and 5 pemetrexed-resistant/SerG6m3-low cell lines for further characterization (**Figure 6D**), and validated their pemetrexed sensitivities using additional dose-response assays (**Figure 6E**). Interestingly, despite the functional link between serine metabolism and nucleotide synthesis, SerG6m3 was not significantly associated with cell proliferation rates across the entire panel or with BrdU staining in the selected cell lines (**Figure S7B,C**). Glucose/glutamine consumption and lactate/glutamate secretion were also no different between the groups (**Figure S7D-G**). To control for the possibility that cell lines with low SerG6m3 have high rates of serine import and/or large intracellular serine pools that reduce labeling from glucose, intracellular and extracellular serine content were measured in selected cell lines. Again, no significant differences were observed (**Figure S7H-J**). These findings link de novo serine biosynthesis to pemetrexed sensitivity in cultured cells, through mechanisms that cannot fully be explained by cell proliferation rates or simple markers of nutrient exchange.

To assess whether SerG6m3 also predicts pemetrexed sensitivity in vivo, three cell lines (two with high and one with low SerG6m3) were implanted into the flanks of nude mice. Pemetrexed treatment was initiated when the mice developed palpable tumors. Pemetrexed caused a marked reduction of tumor growth in both SerG6m3-high cell lines, but the SerG6m3-low cell line was insensitive to pemetrexed in vivo (**Figure 6F-H**).

## Discussion

A growing appreciation of metabolic heterogeneity in cancer cells has increased the need to understand the mechanisms that regulate metabolic pathway preferences and dependencies in tumors. Metabolic heterogeneity presents opportunities to use diverse metabolic features to predict cancer cell behavior, including therapeutic sensitivities. Many relationships between cancer genotypes and metabolism have been established with isogenic systems in which gain or loss of a single mutation is used to identify metabolic differences. These approaches are productive, but they do not account for the impact of genetic heterogeneity (both in the germline and in scores of somatic mutations) on metabolism. Combining even two mutations can dramatically alter the metabolic state compared to either mutation alone. We performed an unbiased assessment of metabolic features in a large panel of highly-annotated cell lines with genetic, histological and therapeutic heterogeneity, all cultured under identical conditions. Therefore, in this study, metabolic features associated with a characteristic of interest (e.g. a single oncogenic mutation) are by definition robust enough to withstand the mitigating effects of private differences.

A few limitations of this study related to the culture conditions warrant mention. We cultured all the cell lines in ambient oxygen conditions in conventional, commercial media with a non-physiological nutrient composition. A few recent studies have emphasized the significant metabolic impact of media formulated to mimic physiological conditions (Cantor et al., 2017; Vande Voorde et al., 2019). We used conventional culture conditions so that we could integrate metabolic phenotyping with orthogonal datasets (transcriptional, proteomic and therapeutic profiling), all of which were obtained under these same conditions. We are encouraged that some of the predictions arising from our phenotyping experiments were validated in vivo, including in patients. But it is likely that similar unbiased studies in physiological media would have added value. We also note that complementing the current dataset with metabolomic profiling would also likely produce a number of interesting associations.

These cell lines were remarkably diverse in the rate of glucose utilization and the ways in which they used glucose. Although the average Lac/Glc ratio was 1.35, this ratio was as low as 0.3, varied 8-fold across the panel, and many cell lines had values less than 1.0. Because this ratio reports the fraction of glucose carbon secreted as lactate, we consider it as a surrogate for the Warburg effect. Thus, the Warburg effect is not a universal feature of these proliferating cancer cell lines. Furthermore, the Lac/Glc ratio correlated only weakly with cell proliferation, and some rapidly proliferating cell lines had low ratios indicating a relatively efficient form of glucose metabolism. As a single variable, glucose consumption did not correlate with the proliferation rate, consistent with the findings of a previous study (Jain et al., 2012). Glucose consumption did correlate very well with glutamine consumption, emphasizing the cooperative utilization of these two major nutrients in cultured cancer cells.

The TCA cycle produces intermediates for energy formation and macromolecular synthesis, thus supporting cell growth through many complementary mechanisms. Several modes of the TCA cycle have been reported in cultured cancer cells (Birsoy et al., 2015; Cheng et al., 2011; DeBerardinis et al., 2007; Le et al., 2012; Metallo et al., 2011; Sullivan et al., 2015; Yuneva et al., 2007). Isotope tracing reports which fuels contribute carbon to the cycle and how these fuels are processed. Despite the consistency of culture conditions throughout our experiments, we observed diverse forms of the TCA cycle in NSCLC cells. All cell lines had extensive anaplerotic input into the TCA cycle, but no single anaplerotic pathway correlated with proliferation rates. As expected, each cell line generally demonstrated blending of multiple forms of the TCA cycle, and it will be interesting to study whether this reflects multiple activities operating concurrently in the same cell or heterogeneity across the population in the dish (e.g. different metabolic activities regulated by cell cycle stage). The most dominant pattern throughout the panel was the use of glucose as *a* precursor for acetyl-CoA and glutamine as *a* precursor for α-ketoglutarate, as indicated by the high CitG6m2 and CitQ6m4 fractions. Pyruvate carboxylation was also detected, and we studied this pathway in greater detail because it is activated in human lung tumors in vivo (Sellers et al., 2015). Isotopologue signatures related to PC activity predicted both PC expression and dependence on PC for colony formation. Reductive metabolism of glutamine, resulting in citrate pools unlabeled by glucose carbon, was surprisingly prominent given the evidence that this pathway is induced by metabolic stressors that suppress acetyl-CoA levels or electron transport chain function, none of which were anticipated in NSCLC cells growing in nutrient-replete conditions. A lack of strong correlations between GDRC labeling and either lactate secretion or the Lac/Glc ratio suggested that GDRC was not stimulated by major defects in oxidative metabolism in these cells.

In isogenic systems, single mutations in oncogenes or tumor suppressors can cause dramatic metabolic perturbations. We set out to identify genotype-metabolism associations robust enough to detect across genetically diverse cell lines, which better reflect the genetic heterogeneity of human cancer. Although we did identify a few such correlations, the overall paucity of associations suggests that cell-autonomous control of cancer cell metabolism arises from the cumulative effects of cell lineage, epigenetic control of gene expression, the constitutional genome and multiple somatically-acquired mutations rather than from single oncogenic drivers. Among the detected associations, cell lines with co-occurring mutations in *KRAS* and *STK11* displayed higher overall labeling in citrate from glucose (i.e. lower CitG6m0) than the rest of the panel. This association was stronger than associations with either mutant *KRAS* (non-site-specific) or mutant *STK11* alone, suggesting metabolic cooperativity between these mutations as suggested elsewhere (Kim et al., 2013; Kim et al., 2017; Liu et al., 2013). In contrast, *EGFR* mutations were associated with GDRC, indicating a different strategy to produce TCA cycle intermediates than in *KRAS*/*STK11* co-mutant cells. We also noted a high content of unlabeled citrate in *EGFR*-mutant tumor tissue from patients with NSCLC, increasing our confidence in the translational potential of the data set. To our knowledge, this is the first example of a correlation between an oncogenic driver and ^13^C labeling features demonstrated to translate from cell culture to human tumors in vivo. Note that none of these patients were infused with ^13^C-glutamine, so it remains to be seen if GDRC is active in *EGFR*-mutant NSCLCs in patients.

In some cases, metabolic features correlated with therapeutic sensitivity. The GDRC phenotype of *EGFR* mutants was accompanied by enhanced sensitivity to EGFR inhibitors, as predicted by the strong epithelial signatures in these cells. Importantly, GDRC and sensitivity to EGFR inhibition appeared to correlate even in cell lines lacking canonical EGFR mutations. This is potentially important because intraoperative isotope infusions in cancer patients may support in situ detection of metabolic phenotypes relevant to therapy. Predicting sensitivity to conventional chemotherapy is particularly challenging, as we generally lack markers to guide safe and effective deployment of these agents. Metabolic phenotyping may provide such markers. A recent study identified an unexpected relationship between sensitivity to the antifolate methotrexate and a metabolic phenotype characterized by histidine catabolism (Kanarek et al., 2018). The latter pathway consumes tetrahydrofolate and renders cells susceptible to the impact of methotrexate on folate pools. We find that sensitivity to pemetrexed, another antifolate, is predicted by de novo serine/glycine synthesis from glucose. This association may arise from the fact that many of the reactions supplied by serine’s contribution to the folate pool are thought to be inhibited by pemetrexed.

Metabolic phenotypes arise from the complex interplay of factors intrinsic and extrinsic to the cancer cell. It is remarkable that the cell lines studied here had such heterogeneous phenotypes despite our use of uniform culture conditions to isolate cell-autonomous regulation of metabolism. Our findings emphasize that diverse metabolic activities can support rapid proliferation of malignant cells. We anticipate that defining cell-autonomous metabolic diversity will contribute to ongoing efforts to understand metabolic phenotypes of human tumors in vivo.

## Supporting information

Merged Supplemental Figures with Legends

## Acknowledgments

We thank Maithili Dalvi, Robin Frink, Rachel Greer and Sunny Zachariah for generating compound sensitivity data for NSCLC cell lines. L.C. was supported by an American Association for Cancer Research (AACR) Basic Cancer Research Fellowship (15 40 01 CAIL). R.J.D. is supported by the Howard Hughes Medical Institute, National Cancer Institute (R35CA22044901), Cancer Prevention and Research Institute of Texas (RP160089), Robert A Welch Foundation (Grant I-1733), Robert L. Moody, Sr. Faculty Scholar Endowment and the Joel B. Steinberg, M.D. Chair in Pediatrics.

## Author Contributions

Conceptualization, R.J.D. and P.H.C.; Methodology, P.H.C.; Formal Analysis, L.C., E.A.M. and H.S.K.; Investigation, P.H.C, C.Y., J.K., B.F., L.B., B.K., J.S. and M.D.S.; Resources, M.P., B.T., I.S., R.B., S.W., K.H., L.G., L.A.B., J.V.H. and J.D.M; Data Curation, L.C. and L.G.; Writing – Original Draft, L.C., P.H.C. and R.J.D.; Writing – Review & Editing L.C. and R.J.D.; Supervision, R.J.D., G.X., J.D.M. and M.A.W.; Funding Acquisition, R.J.D.

## STAR Methods

### KEY RESOURCES TABLE

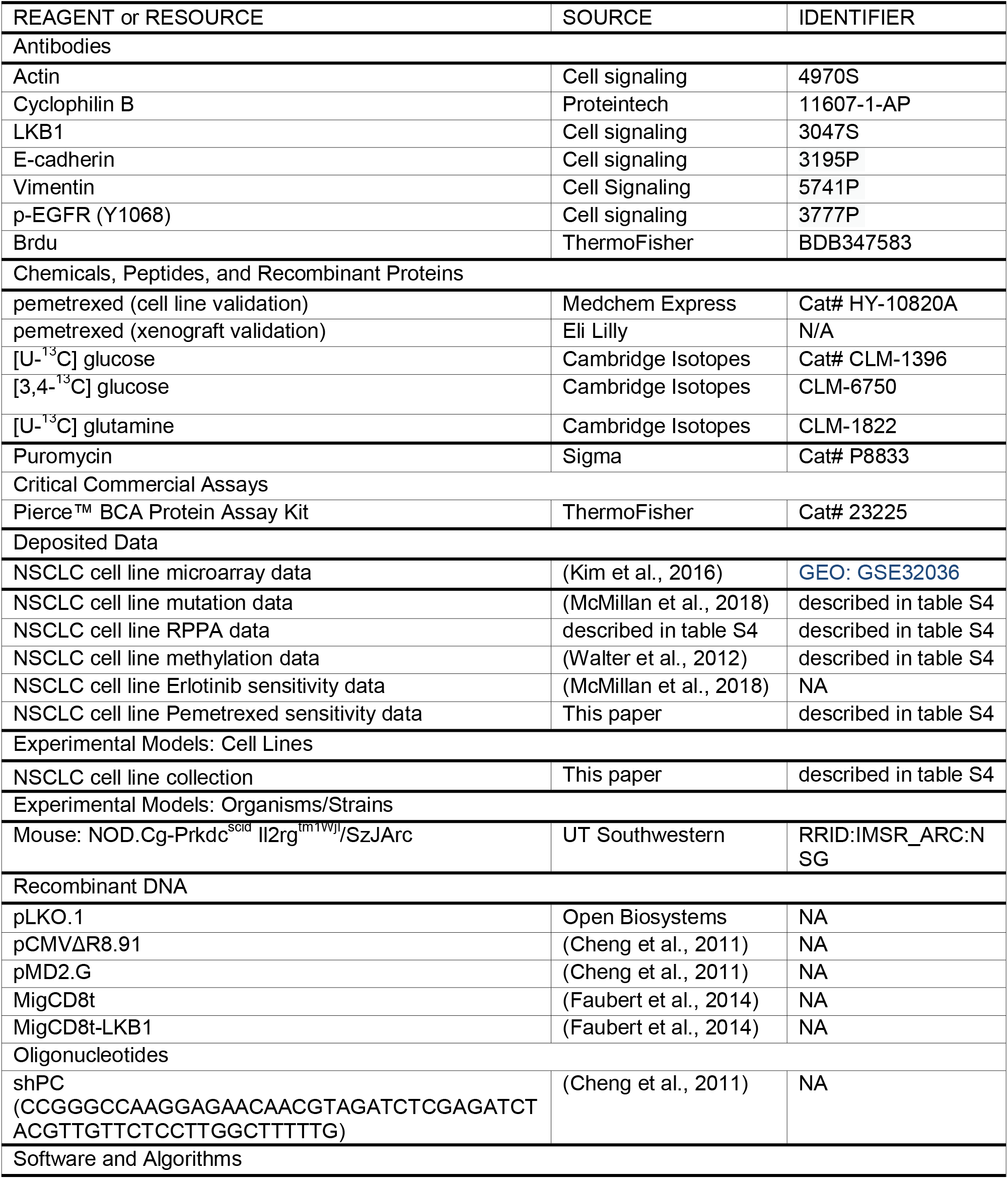
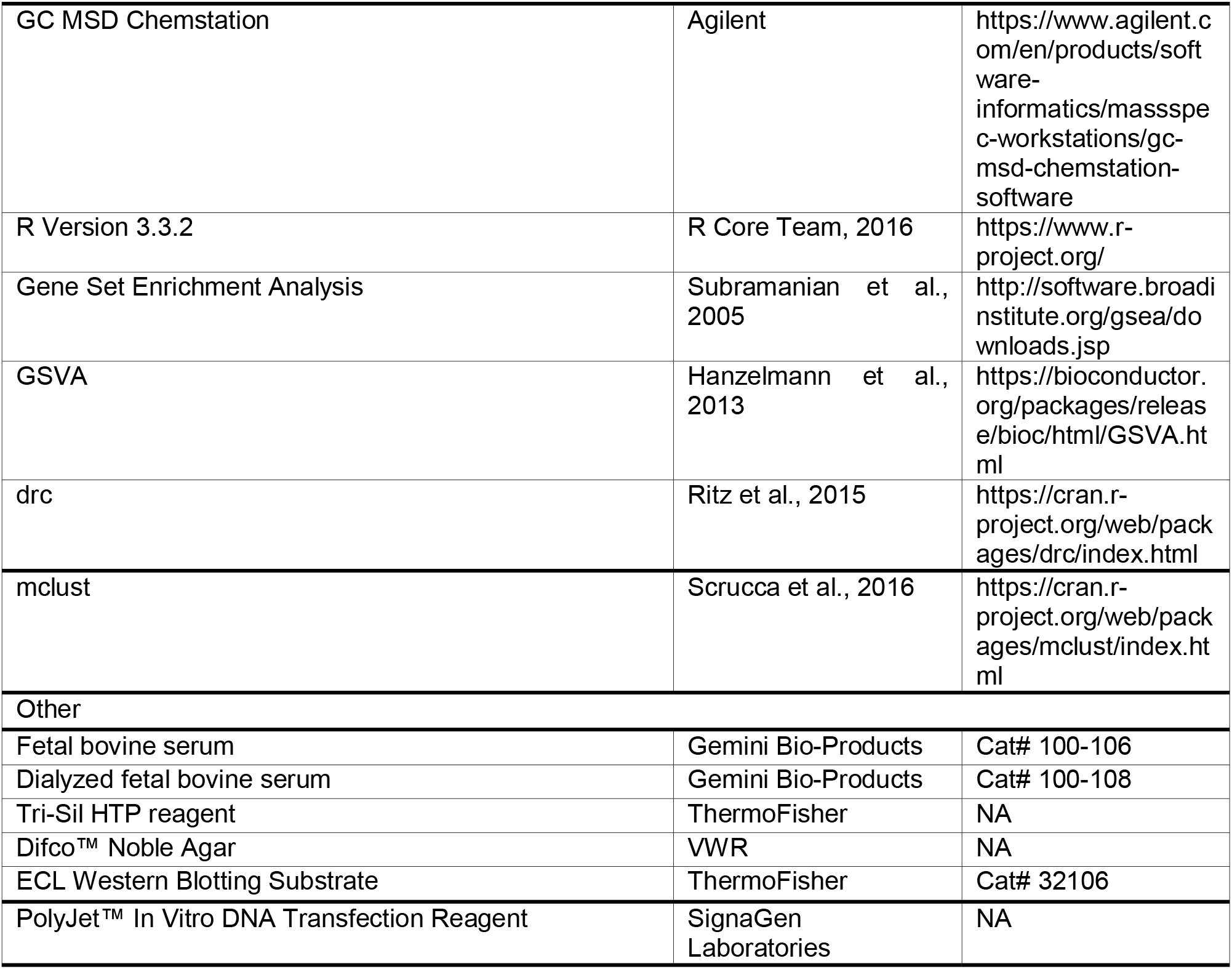

#### Cell culture

Most NSCLC lines used in this study were part of the NCI and HCC (Hamon Cancer Center at UT Southwestern) series, with the exception of A549, Calu-1, Calu-6 (American Type Culture Collection; ATCC), DFCI024, DFCI032 (Dana Farber Cancer Institute, courtesy of Pasi Jänne), PC-9 (Johns Hopkins University School of Medicine, courtesy of Bert Vogelstein). Cell lines were DNA fingerprinted with PowerPlex Fusion24 (Promega). All cell lines were cultured in Roswell Park Memorial Institute medium (RPMI) with 10 mM glucose and 2 mM glutamine supplemented with 5% FBS (Sigma). Nutrient deprivation and metabolic labeling experiments were conducted in RPMI supplemented with 5% dialyzed FBS (Hyclone), sodium bicarbonate (42.5 mM), HEPES (25 mM), Penicillin/Streptomycin (10 U and 10 μg/mL, respectively) and glucose/glutamine as indicated.

#### Stable Isotope labeling analysis

Dishes of 80–90% confluent cells were rinsed twice in PBS, then overlaid with medium containing the isotopically enriched nutrient, and cultured for 6 hours or 24 hours. Note that 4mM glutamine was used because some cell lines depleted the medium of glutamine within 24h if 2mM was used. For analysis of intracellular metabolites by GC/MS, cells labeled in 6-cm dishes were rinsed in ice-cold normal saline and lysed with three freeze-thaw cycles in cold 50% methanol. The lysates were centrifuged to remove precipitated protein, a standard (50 nmols of sodium 2-oxobutyrate) was added, and the samples were evaporated and derivatized by trimethylsilylation (Tri-Sil HTP reagent, Thermo Scientific). Three microliters of the derivatized material was injected onto an Agilent 6970 gas chromatograph equipped with a fused silica capillary GC column and networked to an Agilent 5973 mass selective detector. Retention times of all metabolites of interest were validated using pure standards. The abundance of the following ions was monitored: m/z 245-249 for fumarate, m/z 335-339 for malate, m/z 219-222 for lactate, m/z 306-309 for serine, m/z 276-278 for glycine, and m/z 465–471 for citrate. The measured distribution of mass isotopologues was corrected for natural abundance of ^13^C.

#### Nutrient utilization rates

To measure metabolic rates, one million cells were plated into 6-cm dishes and cultured until 90% confluent. At time 0, the cells were rinsed in PBS, fed with 1.5 mL of RPMI with 10 mM glucose, 2 mM glutamine and dialyzed FBS, and cultured. End-point experiments proceeded for 7 hours, then the medium was collected and analyzed for metabolite abundance. Concentrations of glucose, lactate, glutamine, and glutamate were determined from 0.6-mL aliquots of medium using an automated electrochemical analyzer (BioProfile Basic-4 analyzer; NOVA). Metabolic rates were determined by normalizing absolute changes in metabolite abundances to protein content.

#### Cell growth and nutrient dependence assay

To monitor proliferation, cells were seeded at 5,000/well in 48-well plates. The next day, cells were replenished with 0.5 ml of test medium (complete RPMI, RPMI without glucose, or RPMI without glutamine). After 1, 3 and 5 days, DNA content was determined by adding 0.25 ml water to each well and freezing at −80°C for 2 hr. The cells were then warmed to room temperature, and 0.5 ml of 0.1 μg/ml Hoechst 33258 in TNE buffer (2 M NaCl, 10 mM Tris-HCl [pH 7.4], and 1 mM EDTA) was added. The plate was incubated in the dark at room temperature, and O.D. at 350 nm was measured using a plate reader. Cell proliferation and survival measurements were made by normalizing DNA content of cells from Day 3 and Day 5 to that of Day 1.

#### Protein expression

Whole-cell lysates were prepared in RIPA buffer and quantified using the BCA protein assay (Thermo Scientific). Protein was separated by SDS/PAGE and transferred to a PVDF membrane. The membrane was blocked overnight at 4°C in PBS with Tween 20 (PBST) containing 5% milk, then probed with primary antibodies overnight at 4°C.

#### LKB1 over-expression

The MigCD8t control and LKB1 overexpression plasmids were kindly provided by Dr. Russell Jones (Van Andel Research Institute). Selected cell lines were infected with supernatants containing MigCD8t or MigCD8t-LKB1 retrovirus. Three days later, flow cytometry was used to enrich for infected cells expressing the CD8t surface marker (Nicolas Loof, the Moody Foundation Flow Cytometry Facility of UTSW).

#### RNA Interference

shPC was cloned into pLKO.1 backbone purchased from Open Biosystems. Lentiviral particles were produced by co-transfecting 293FT cells with the lentiviral construct, pCMVΔR8.91, and pMD2.G using polyjet (SignaGen Laboratories). Virus-containing supernatant was collected 2 days after the transfection and used to infect cells. Puromycin (1 μg/mL) was added 2 days after infection, and selection was continued for 7 days before any experiments.

#### Soft agar colony formation assay

1000 cells were plated into 12-well plates with 0.33% Noble agar (Difco). After 2 weeks, the cultures were stained with 0.05% crystal violet in 20% methanol. Colonies >200 μm in diameter were counted.

#### BrdU staining

Cells were cultured with 10uM BrdU for one hour and fixed in 70% ethanol at −20°C. After washing with phosphate/citric buffer (40ml Na2HPO4 with 4ml 0.1M citric acid), the cells were incubated with anti-BrdU antibody for one hour, then stained with propidium iodide (PI). For PI/RNase staining, cells were incubated with 0.5 mL of PI/RNAse staining buffer (BD Pharmingen) at 25°C for 15 minutes, then analyzed by flow cytometry.

#### Free amino acid concentration quantitation

Consumption and secretion of amino acids were measured by HPLC (Hitachi; L8900). One million cells were cultured in 6cm dishes and switched to RPMI medium with dialyzed FBS for 7 hours. Medium samples from before and after culture were combined 1:1 with 1M sulfosalicylic acid, followed by centrifugation at 10000 rpm for 15 min at 4°C. The supernatant was then combined with an AEC standard (1:10 by volume) and analyzed by HPLC.

#### Xenograft studies

Female *NOD*/*SCID* mice were obtained from Wakeland laboratories, UT Southwestern Mouse Breeding Core (Dallas, TX) at approximately 6-8 weeks of age. 1×10^6^ cancer cells were injected subcutaneously into the shaved right flank and monitored by calipers (v = π/6*|*w^2^; v, volume; l, length, w, width). All mouse procedures were performed in compliance with UT Southwestern IACUC policies.

#### Gene set enrichment analysis (GSEA) and single sample GSEA (ssGSEA)

The C2 (curated gene sets) library was downloaded from Molecular Signatures Database (MSigDB). GSEA was performed in continuous mode with the Spearman rank correlation as the ranking metric. The analysis used an R script adapted from R-GSEA. Scores from ssGSEA were calculated by implementing the “gsva” function from the R package “GSVA” (Hanzelmann et al., 2013).

#### Pemetrexed sensitivity assay

Selected cell lines were treated with different doses (0.01, 0.05, 0.1 and 1 μM) of pemetrexed for 3 days. A standard 4-parameter log-logistic fit between the survival rate and the dosage was generated by the “drm” function from the R package “drc” (Ritz et al., 2015). Areas under the fitted dose response curve were calculated and compared by t-test.

#### Stepwise regression

Feature selection for the Erlotinib sensitivity prediction model was performed by stepwise regression using “stepAIC” function with “both,” as directed by the R package “MASS”. Coefficient tables from multiple regression models were visualized with the “tab_model” function from the R package “sjPlot”.

