## Supplementary material for "Metabolic Diversity in Human Non-Small Cell Lung Cancer Cells": Merged Supplemental Figures with Legends

**Legends for Supplementary Figures**

**Figure S1. Low levels of variation among biological replicates and high levels of cell-autonomous metabolic diversity among cell lines.**

(**A**). Scatterplot comparing the averaged variations among biological replicates for the same cell line (x-axis values) and variations among all cell lines (y-axis values) for isotopologue enrichment data from $^{13}C$ tracing experiments. For each isotopologue, the color represents the metabolite of interest, shape represents the tracer, and transparency represents the labeling time. Standard deviations were calculated as variation estimates; the standard deviations of the between-replicate standard deviations were also calculated and plotted as horizontal tails extending to the right for each isotopologue.

(**B**). Mass isotope distribution (MID) of citrate labeled by [U-$^{13}C$]glucose for 6 hours in biological replicates of 3 cell lines. For each cell line, the replicate experiments were conducted with gaps of several months in between.

(**C** and **D**). Distinct MID pattern of citrate for different cell lines, labeled by [U-$^{13}C$]glucose (**C**) or [U-$^{13}C$]glutamine (**D**) for 6hr, indicating the striking diversity of labeling patterns among these cell lines.

**Figure S2. Mass isotopologue distribution and variation for selected metabolites from [U-$^{13}C$]glucose or [U-$^{13}C$]glutamine labeling.**

(**A**-**E**). Violin plots showing mass isotopologue distribution in fumarate (**A**), malate (**B**), lactate (**C**), serine (**D**) and glycine (**E**).

(**F**). Variance of different isotopologues.

**Figure S3. Correlation heatmap displaying pairwise Pearson correlations among all metabolic features**

These correlations are based on isotopologue fractions from $^{13}C$ tracing experiments at 6h and at 24h as well as features from nutrient utilization, cell growth and nutrient dependency. Larger sizes and darker shades of the circles reflect more significant correlations. Red circles indicate positive correlations and blue circles indicate negative correlations.

**Figure S4. Estimation of pyruvate carboxylation dependent anaplerosis after [U-$^{13}C$]glucose labeling**

(**A**). Malate m+3 can be generated from [U-$^{13}C$]glucose after two rounds of the TCA cycle. This confounds the interpretation of Malate m+3 as an indicator of pyruvate carboxylase activity. Citrate m+4 is the nearest detectable TCA cycle intermediate in this route that cannot be reversibly generated from malate m+3. Therefore it might serve as a predictor for Malate m+3 generated through this pathway.

(**B**). Residuals from regressing MalG6m3 on CitG6m4 might represent MalG6m3 produced independently from the pathway in (**A**), the majority of which likely arises from pyruvate carboxylation.

(**C**-**D**). Scatter plot showing correlations between *PC* gene expression and MalG6m3 (**C**) or residuals from regressing MalG6m3 on CitG6m4 (**D**). The correlation coefficient and p-value in the title are based on Pearson correlation.

**Figure S5. Re-expression of LKB1 in three selected cell lines with concurrent mutations in *KRAS* and *STK11*.**

Western blot showing LKB1 levels in parental and re-expression cell lines.

**Figure S6. The GDRC phenotype is associated with epithelial state**

(**A**-**B**). Gene Set Enrichment Analysis (GSEA) identified CitQ6m5 is positively correlated with ZEB1 target genes (**A**) and negatively correlated with Gefitinib resistance genes (**B**).

(**C**). CitG6m0 and CitQ6m5 are significantly different between the epithelial and mesenchymal groups, whereas similar levels of Lac/Glc and Glu/Gln were observed in the two groups. Epithelial and mesenchymal group classification was based on hierarchical clustering with the EMT gene expression signature in Figure 5D.

(**D**). Scatterplot and pairwise Pearson correlations among *CDH1* methylation, *CDH1* gene expression, *CDH1*-encoded E-cadherin expression and CitG6m0. ***, p<=0.001; **, p<=0.01; *, p<=0.05; no asterisk is used when p>0.05.

**Figure S7. Association between SerG6m3 and serine metabolism genes and comparison of various metabolic features in selected SerG6m3-high and -low cells**

(**A**). Heatmap showing expression of selected genes involved in serine metabolism. Correlation coefficients and p-values from Spearman correlations between gene expression and SerG6m3 are provided.

(**B**). Non-significant correlation between SerG6m3 and cell growth feature Day3/Day1.

(**C-J**). Insignificant results from comparing Brdu staining (**C**), glucose uptake (**D**), lactate secretion (**E**), glutamine uptake (**F**), glutamate secretion (**G**), intracellular serine pool size inferred from GC-MS (**H**), intracellular (**I**) or extracellular (**J**) serine concentrations measured by HPLC in SerG6m3-high and -low cell lines.

### Figure S1

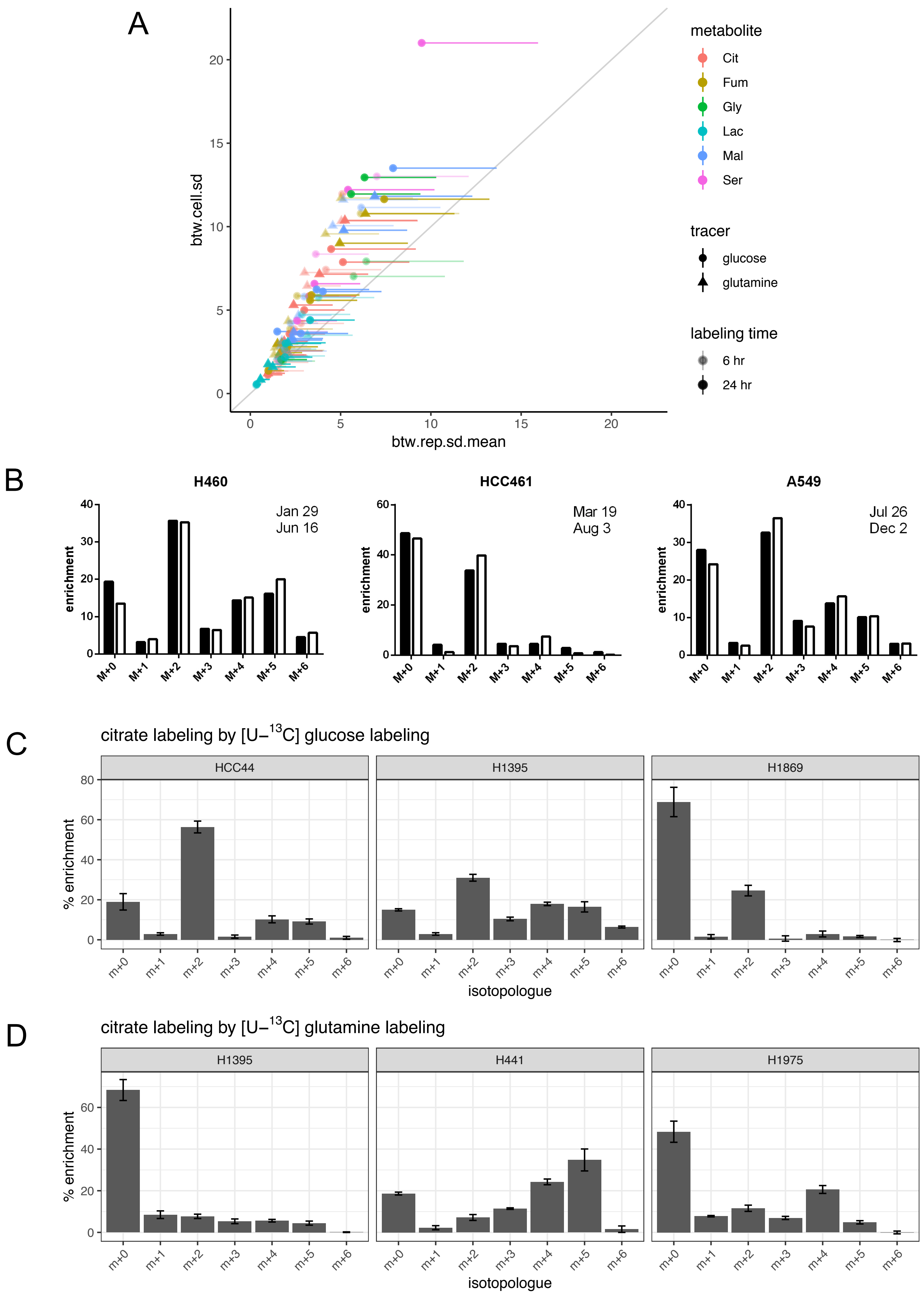
A
btw.cell.sd
btw.rep.sd.mean
metabolite
Cit
Fum
Gly
Lac
Mal
Ser
tracer
glucose
glutamine
labeling time
6 hr
24 hr
B
H460
Jan 29
Jun 16
HCC461
Mar 19
Aug 3
A549
Jul 26
Dec 2
enrichment
M+0
M+1
M+2
M+3
M+4
M+5
M+6
C
citrate labeling by [U–13C] glucose labeling
HCC44
H1395
H1869
% enrichment
isotopologue
D
citrate labeling by [U–13C] glutamine labeling
H1395
H441
H1975
% enrichment
m+0
m+1
m+2
m+3
m+4
m+5
m+6
isotopologue

### Figure S2

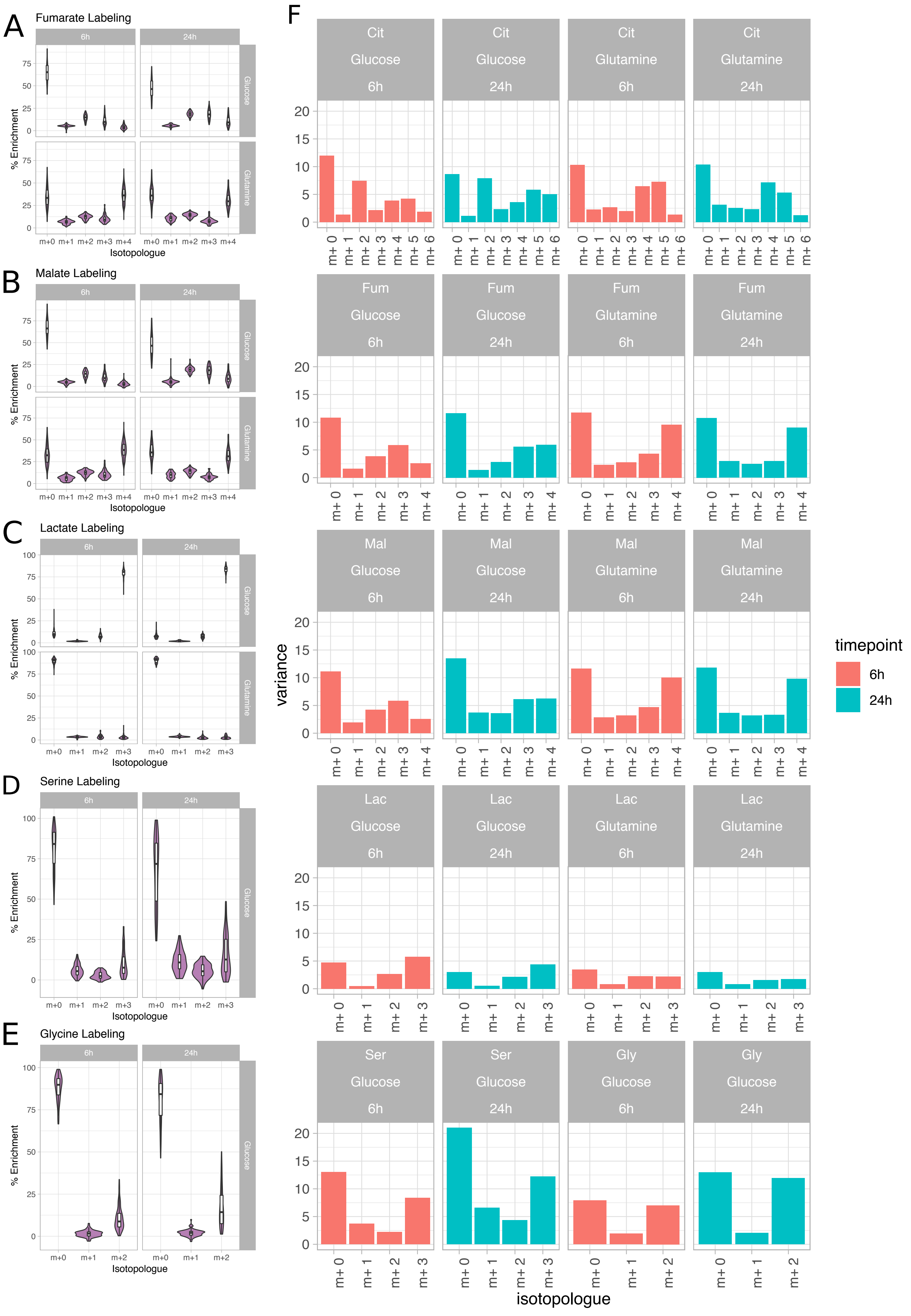
A
Fumarate Labeling
B
Malate Labeling
C
Lactate Labeling
D
Serine Labeling
E
Glycine Labeling
6h
24h
Glucose
Glutamine
% Enrichment
Isotopologue
m+0 m+1 m+2 m+3 m+4
F
Cit Glucose 6h
Cit Glucose 24h
Cit Glutamine 6h
Cit Glutamine 24h
Fum Glucose 6h
Fum Glucose 24h
Fum Glutamine 6h
Fum Glutamine 24h
Mal Glucose 6h
Mal Glucose 24h
Mal Glutamine 6h
Mal Glutamine 24h
Lac Glucose 6h
Lac Glucose 24h
Lac Glutamine 6h
Lac Glutamine 24h
Ser Glucose 6h
Ser Glucose 24h
Gly Glucose 6h
Gly Glucose 24h
variance
isotopologue
timepoint
6h
24h

### Figure S3

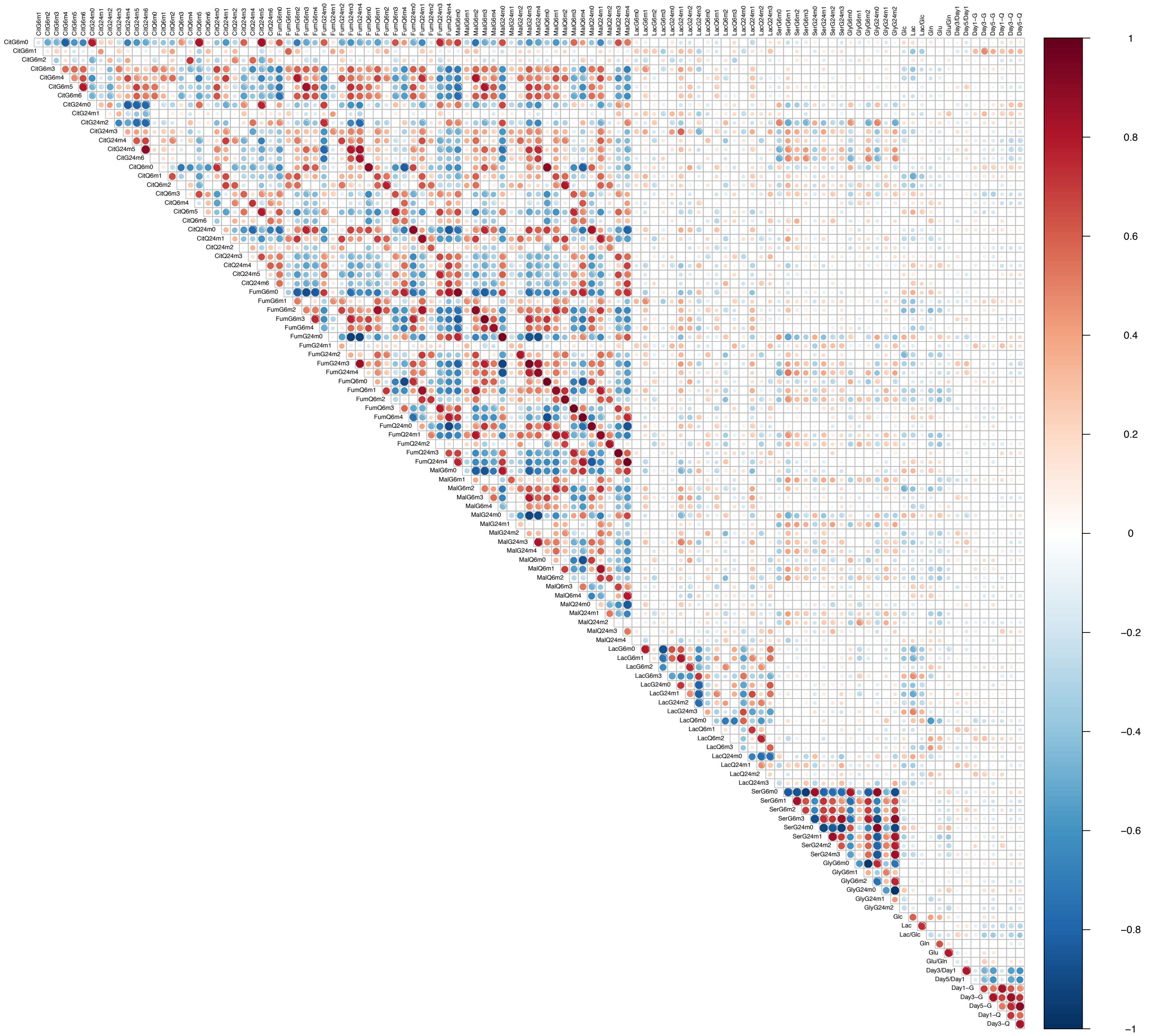

### Figure S4

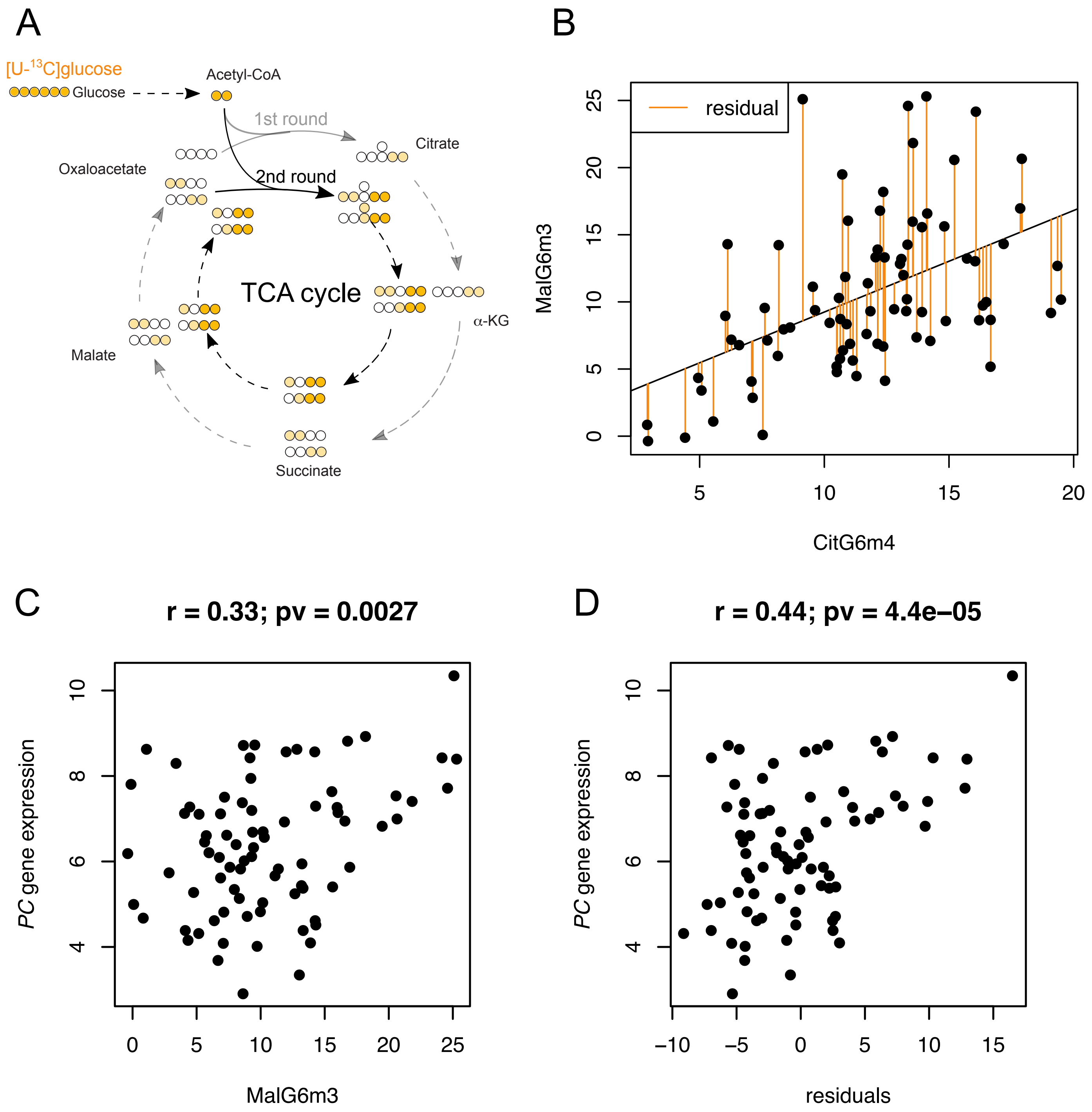
A
[U-13C]glucose
Glucose
Acetyl-CoA
1st round
2nd round
Citrate
Oxaloacetate
TCA cycle
α-KG
Malate
Succinate
B
residual
MalG6m3
CitG6m4
C
r = 0.33; pv = 0.0027
PC gene expression
MalG6m3
D
r = 0.44; pv = 4.4e−05
PC gene expression
residuals

### Figure S5

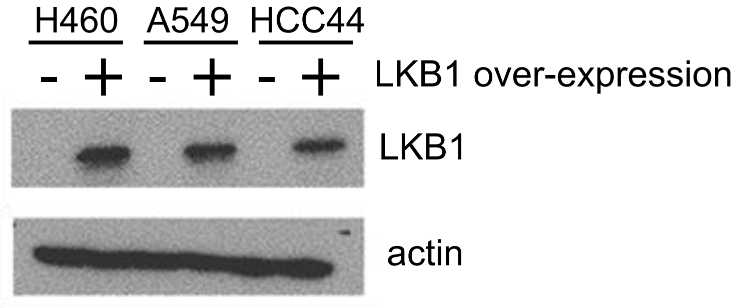

### Figure S6

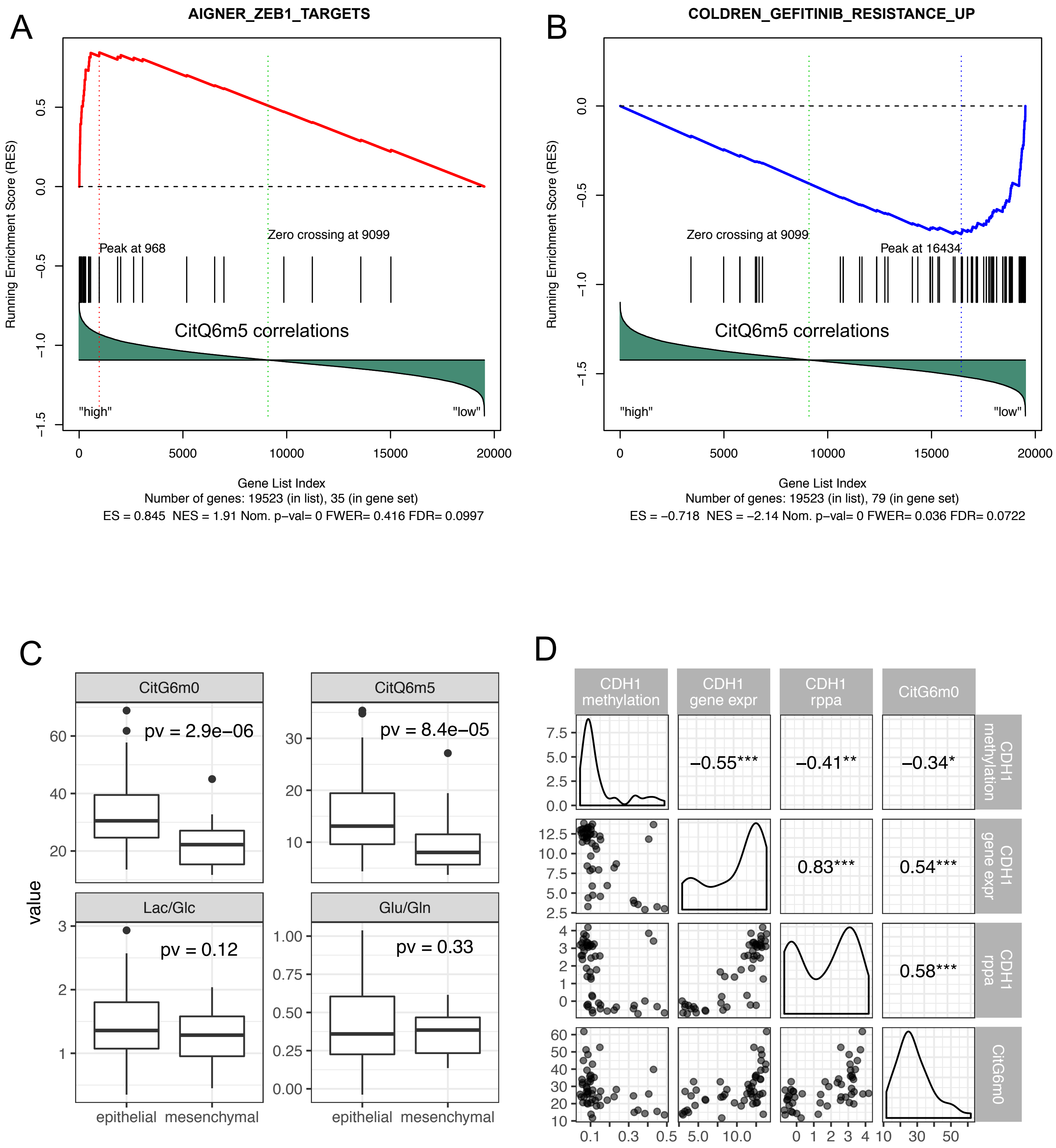
A
AIGNER_ZEB1_TARGETS
Running Enrichment Score (RES)
Peak at 968
Zero crossing at 9099
CitQ6m5 correlations
"high"
"low"
Gene List Index
Number of genes: 19523 (in list), 35 (in gene set)
ES = 0.845 NES = 1.91 Nom. p-val= 0 FWER= 0.416 FDR= 0.0997
B
COLDREN_GEFITINIB_RESISTANCE_UP
Running Enrichment Score (RES)
Zero crossing at 9099
Peak at 16434
CitQ6m5 correlations
"high"
"low"
Gene List Index
Number of genes: 19523 (in list), 79 (in gene set)
ES = −0.718 NES = −2.14 Nom. p-val= 0 FWER= 0.036 FDR= 0.0722
C
value
CitG6m0
pv = 2.9e−06
CitQ6m5
pv = 8.4e−05
Lac/Glc
pv = 0.12
Glu/Gln
pv = 0.33
epithelial mesenchymal
epithelial mesenchymal
D
CDH1 methylation
CDH1 gene expr
CDH1 rppa
CitG6m0
−0.55***
−0.41**
−0.34*
0.83***
0.54***
0.58***

### Figure S7

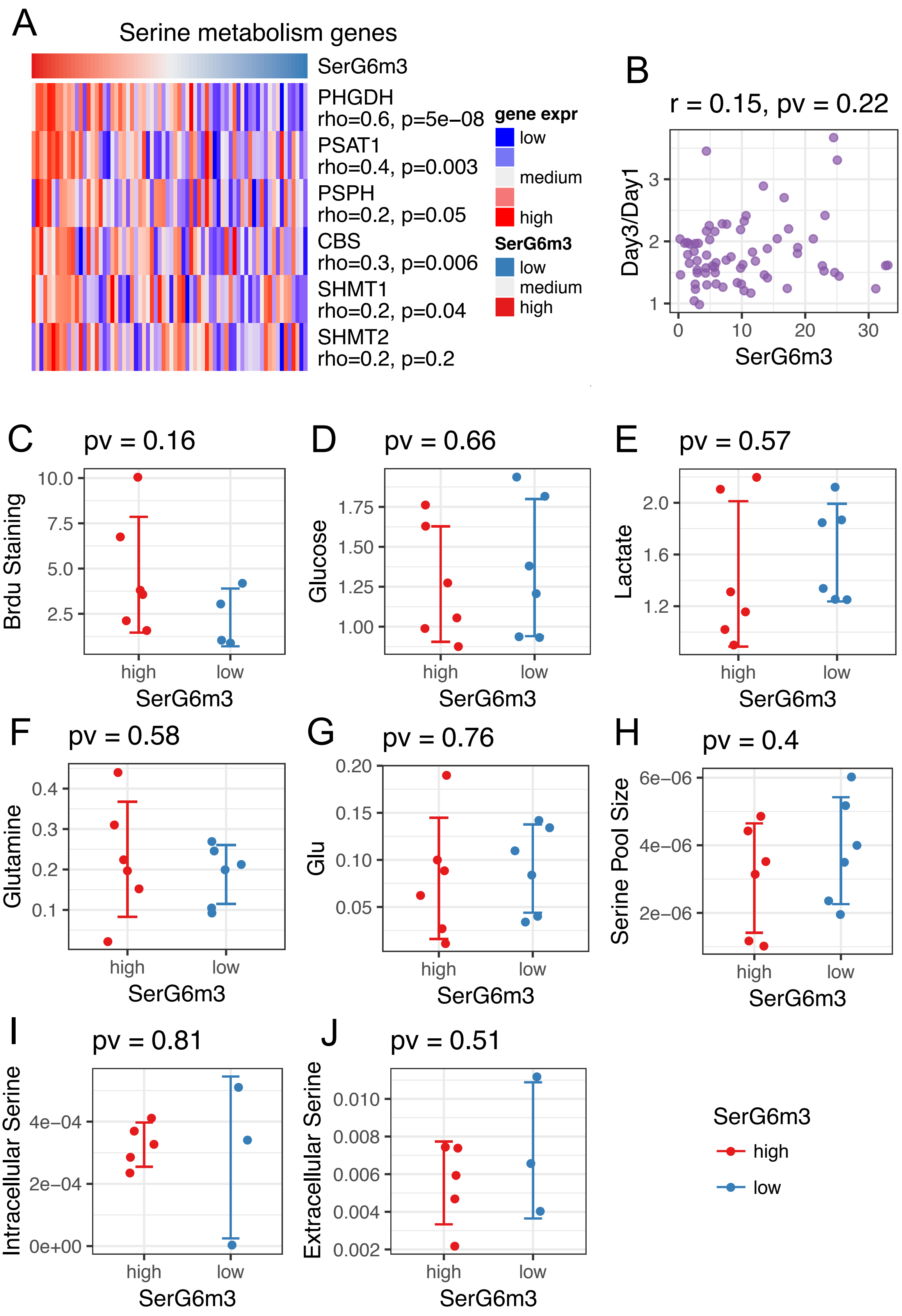

A
Serine metabolism genes
SerG6m3
PHGDH
rho=0.6, p=5e−08
PSAT1
rho=0.4, p=0.003
PSPH
rho=0.2, p=0.05
CBS
rho=0.3, p=0.006
SHMT1
rho=0.2, p=0.04
SHMT2
rho=0.2, p=0.2
gene expr
low
medium
high
SerG6m3
low
medium
high
B
r = 0.15, pv = 0.22
Day3/Day1
SerG6m3
C
pv = 0.16
Brdu Staining
D
pv = 0.66
Glucose
E
pv = 0.57
Lactate
F
pv = 0.58
Glutamine
G
pv = 0.76
Glu
H
pv = 0.4
Serine Pool Size
I
pv = 0.81
Intracellular Serine
J
pv = 0.51
Extracellular Serine
high
low
SerG6m3
